# Multimodal Sensitivity Enhancement of Binding-based Diagnostic Assays Using Functional Hydrogels

**DOI:** 10.64898/2026.09.21.753162

**Authors:** Pranshu Rajurkar, Samatha de Gorostiza Wise, Aniruddh Sarkar

**Affiliations:** Wallace H Coulter Department of Biomedical Engineering, Georgia Institute of Technology and Emory University School of Medicine, Atlanta GA 30332

## Abstract

Immunoassays are critical for clinical diagnostics, yet their performance is restricted by the physical constraints of flat surfaces used to immobilize capture molecules which limit loading capacity and induce protein denaturation. Hydrogels provide immense volumetric molecular loading capacity but translating them across diverse assay formats and readout modalities without compromising analyte permeability remains challenging. Here, we present a universal strategy for hydrogel integration into immunoassay platforms for enhanced optical and electronic biosensing. Utilizing bio-orthogonal photo-click chemistry, we developed a modular and tuneable poly(ethylene glycol) (PEG) matrix that achieves a 6,000-fold increase in molecular loading and a large mesh size for analyte diffusion. This 3D architecture demonstrates broad multimodal utility with easy integration into a range of binding-based assays enabling higher sensitivity, repeatability and multiplexability. It achieves high sensitivity enhancement for fluorescence detection in microarrays (38-fold) and microtiter plates (25-fold) and enables multiplexed readout in single wells, including from clinical serum samples. In silver metallization-based assays it boosts densitometric optical detection sensitivity (23-fold) and enables inexpensive electronic detection. In lateral flow assays, it generates 20-fold signal enhancement while enabling multiplexability and higher repeatability. By standardizing high-capacity volumetric biosensing, this scalable, plug-and-play technology empowers next generation highly sensitive, field-deployable point-of-care diagnostics.

## 1. Introduction

Diagnosis of diseases is commonly achieved by quantifying specific biomarkers present in complex biological matrices such as serum using immunoassays. The fundamental principle of immunosensing relies on the specific affinity interactions between a capture molecule (CM) and its cognate target analyte. Leveraging these interactions, binding-based immunoassays – including enzyme-linked immunosorbent assays (ELISAs), microarrays, and lateral flow assays (LFAs) – have become indispensable tools in modern clinical diagnostics, biomedical research and biosensor development^1^. By converting molecular recognition events into detectable optical or electronic signals, these platforms provide the information necessary for biomarker discovery, disease diagnostics, monitoring therapeutic efficacy, and conducting epidemiological surveillance. Different immunoassay modalities are optimized and used in different contexts such as in centralized lab-based clinical diagnostics (ELISAs), high-throughput biomarker discovery and research (microarrays) and inexpensive diagnostics for point-of-care or home use (LFAs).

The analytical performance of these immunoassays is governed by the efficiency with which CMs are immobilized on the detection substrate^2^. Traditionally, these assays rely on the passive adsorption of CMs onto 2D planar substrates. Driven by non-specific hydrophobic and electrostatic interactions, this process often introduces significant biophysical constraints. Direct contact with rigid, hydrophobic surfaces can induce protein denaturation, while random molecular orientation can mask up to 95% of active binding sites, rendering them inaccessible to the target analyte^3^. Critically, the limited surface area of 2D substrates imposes a strict ceiling on loading capacity, which, coupled with high background noise from non-specific matrix adsorption, severely restricts the sensitivity and dynamic range of the platform^4^. While developing higher-affinity binding pairs or advanced detection techniques can improve assay sensitivity, these approaches often introduce significant costs, complex instrumentation, and scalability hurdles^5,6^. A compelling alternative is altering the sensing substrate itself to optimize CM presentation. Although chemically functionalized surfaces and pseudo-3D architectures like polymer brushes or dendrimers increase CM loading and improve orientation, they remain fundamentally limited by spatial crowding^7–10^. Highly dense monolayers and tightly packed macromolecular chains can impede the accessibility of larger target analytes, induce protein denaturation, and promote non-specific interactions due to their proximity to the rigid solid substrate^11^.

A more robust solution to these interfacial challenges has emerged through the development of three-dimensional (3D) hydrogels as scaffolds for immobilizing CMs^12–14^. Unlike planar or pseudo-3D substrates, hydrogels provide a highly hydrated, biomimetic environment that effectively preserves the native conformation and biological activity of immobilized CMs^15^. This transition from a surface-bound model to a volumetric framework facilitates significantly higher loading densities of biorecognition elements and steric freedom for targets to approach from all directions^16,17^. Furthermore, the crosslinked architecture of a hydrogel offers tuneable porosity, which can be engineered to enable the facile diffusion of target analytes while providing size-based exclusion of larger interferents that may cause non-specific binding^18,19^. Poly(ethylene glycol) (PEG)-based hydrogels are particularly advantageous in this context as their inert polymer backbone minimizes fouling in complex clinical matrices, while their chemical modularity allows for precise control over the structural and functionalization properties of the network^20,21^. To this end, PEG-based hydrogels have been previously explored to enhance specific biosensing modalities. In fluorescence-based microarrays, functionalized 3D scaffolds have been developed to increase CM density and preserve native protein conformation, outperforming traditional 2D planar slides^22^. Similarly, the high-sensitivity readout of enzymatic silver metallization that enables a simple, densitometric optical quantification of analyte binding has been adapted for hydrogel platforms^23,24^. Furthermore, in the context of LFAs, hydrogels have been incorporated as flow-control traps on paper substrates to slow down wicking rates, extend target incubation times and thus amplify signal readouts^25,26^.

Despite these modality-specific advancements, a fundamental challenge in developing 3D immobilization matrices remains managing the inherent trade-off between functional loading and matrix permeability. Increasing CM density can inadvertently diminish hydrogel porosity and impede target diffusion, rendering the bulk of the scaffold inaccessible to target analytes^22^. To navigate this biophysical hurdle, current 3D scaffolds are typically meticulously optimized for each single assay format and readout technique. Consequently, the field relies on a fragmented variety of formulations that are difficult to translate between sensing modalities, lacking a unified chemical architecture. This lack of standardization contributes to the lack of wide use of hydrogels in immunoassays for clinical diagnostics or research despite the above demonstrated use cases. Furthermore, their potential as functional scaffolds for emerging electronic transduction techniques remains largely untapped. Similarly, their capacity to enhance CM immobilization on paper-based substrates in LFAs is still largely unexplored. Ultimately, there is a clear need for a versatile, scalable interface that maintains both high volumetric capacity and efficient target transport regardless of the specific diagnostic modality.

To address the need for a unified and scalable sensing interface, we present a versatile, ‘plug-and-play’ hydrogel platform based on thiol-norbornene photo-click chemistry^27^ as a standardized functional immobilization scaffold for high-performance diagnostics. This PEG-based framework utilizes a biorthogonal, stoichiometric reaction that allows for temporal control over crosslinking and facile covalent functionalization with thiolated CMs^28^. By achieving a unique synergy between high functionalization density and a large mesh size, this architecture creates an optimal 3D environment that bypasses the surface-area constraints and steric hindrance of planar substrates. We demonstrate the broad utility of this platform by achieving significant analytical enhancements across multiple disparate sensing modalities including fluorescence-based microarrays, silver metallization-enabled optical and electronic readouts, and LFAs. Furthermore, we illustrate that the modularity of this formulation enhances multiplexing capabilities in both fluorescence and LFA formats. This study establishes a common chemical architecture that is easily integrated and scalable, providing a generalizable yet versatile interface that transforms how CMs are immobilized across diverse diagnostic platforms regardless of the sensing modalities.

## 2. Results and Discussion

### 2.1. Analyzing the Effect of Increasing CM Density on Target Binding

The transition from a 2D surface-bound model to a 3D volumetric framework enables a substantial increase in the loading capacity of CMs **(Figure 1A)**. Traditionally for assays for low abundance analytes, it is often assumed that the CMs on a 2D surface are already in vast excess and thus not the limiting component of the system. Here, we sought to use a simplified analytical framework to investigate the impact of increasing CM density on analyte capture and detection sensitivity.

**Figure 1.**
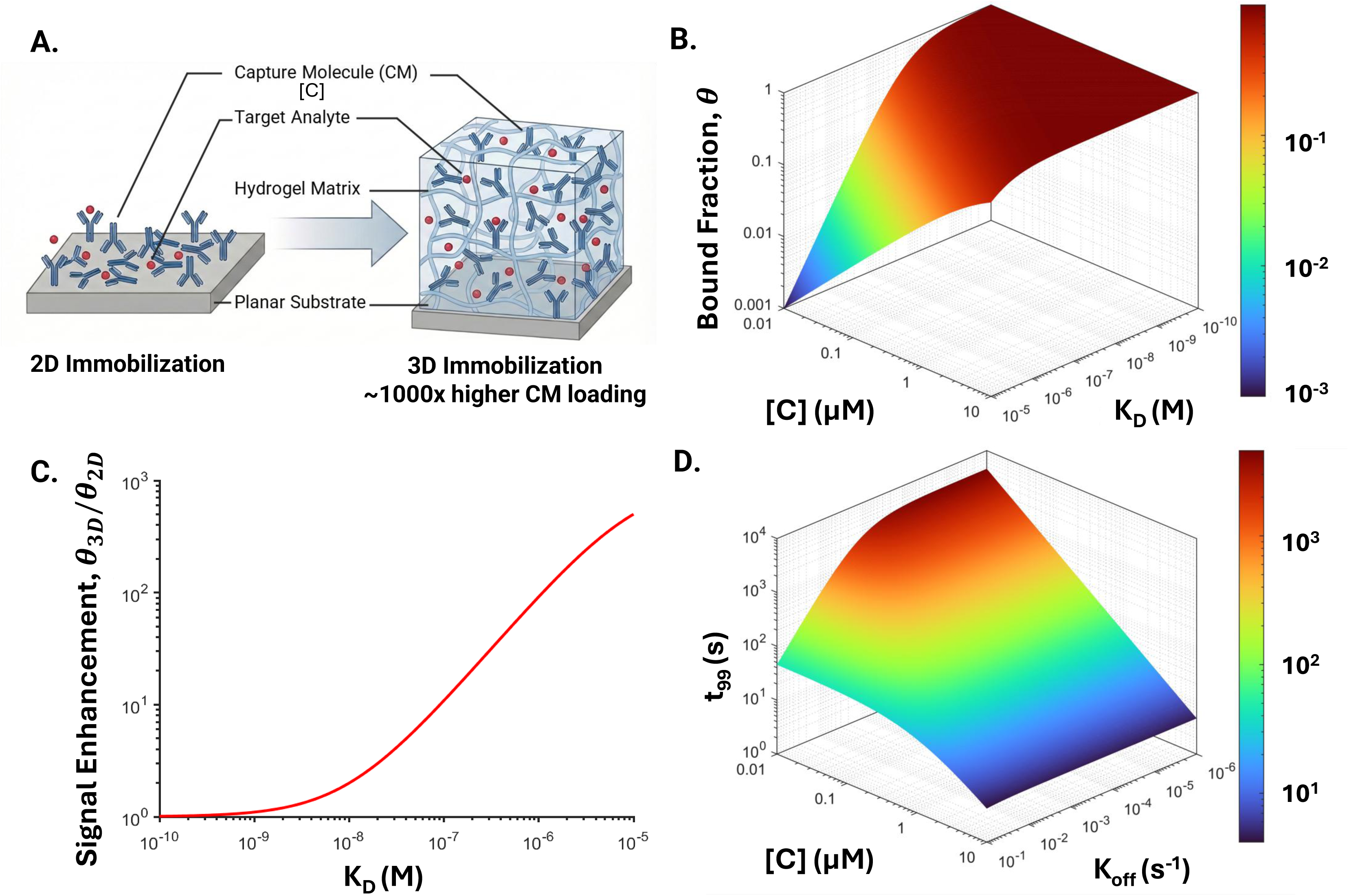
Theoretical modeling of target binding equilibrium and kinetics in 2D planar and 3D volumetric platforms. **(A)** Schematic representation comparing the theoretical capture molecule loading capacity of a 2D surface-bound planar model to a 3D volumetric framework. **(B)** 3D plot modeling the fraction of target bound at equilibrium (θ_T_) as a function of capture molecule concentration ([A]) and equilibrium dissociation constant (K_D_). **(C)** Theoretical signal enhancement (fold change) provided by the 3D volumetric framework over the 2D surface **(D)** 3D kinetic plot modeling the time required to reach 99% equilibrium (t_99_) as a function of the k_off_ and [A], evaluated at a constant k_on_ of 10^5^ M^−1^s^−1^

For simplicity, we consider here a bimolecular binding reaction:

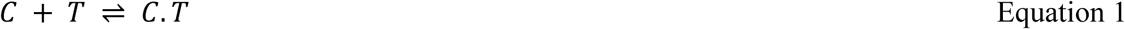

where C is the CM, T is the target analyte and C.T is their binding product. To establish a theoretical baseline for comparison, we first equated surface-bound CM density to an effective bulk concentration [*C*] relative to a standard 100 μL sample volume. For example, for a typical high-binding 96-well plate, with a maximum IgG antibody loading^3^ of 500 ng cm^-2^, commonly used for ELISAs, this yields the effective concentration of approximately 10 nM. Based on prior hydrogel work, we also assumed a conservative 1,000-fold enhancement in loading^19^, yielding an effective enhanced CM concentration of 10 μM.

Under the typical condition that the CM is in vast excess ([*C*] ≫ [*T*]), the fraction of target bound at equilibrium (*θ_T_*) can be derived from the Langmuir isotherm as:

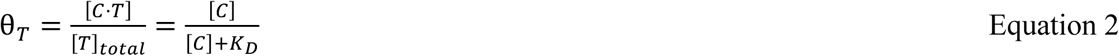

where *K_D_* is the equilibrium dissociation constant. This bound fraction is plotted as a function of [*C*] & *K_D_* in **Figure 1B**. This plot highlights that for high-affinity binders (*K_D_* < 100 pM) near-complete target capture is achieved even at 2D surface densities ([*C*] ≈ 10 *nM*). However, for low-to-moderate affinity interactions, increases in [*C*], as can be achieved in 3D volumetric frameworks, drastically improves equilibrium target binding. For instance, at a *K_D_* of 10 μM, a traditional 2D surface captures less than 0.1% of the target, effectively yielding no signal. In contrast, the 3D scaffold ([*C*] ≈ 10 *μM*) shifts the equilibrium to capture 50% of the available analyte, enabling up to a 500-fold signal enhancement **(Figure 1C)**. This demonstrates that the 3D volumetric framework expands the repertoire of usable biorecognition elements to include those with low target affinities (e.g. antibodies for novel targets such as from emerging pathogens, aptamers for small molecules, lectins for glycans etc.) enabling high-performance sensing even with suboptimal binder affinity.

Furthermore, the volumetric excess of [*C*] significantly accelerates the binding kinetics. The time required to reach 99% equilibrium (*t*_99_) is dictated by the following equation^4^:

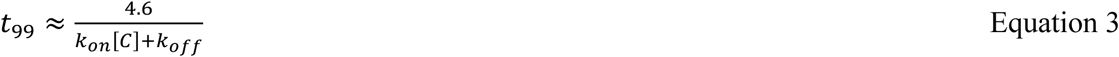

Using an association rate constant (*k_on_*) of a typical value of 10^5^ M^-1^ s^-1^, this kinetic model highlights a significant kinetic bottleneck for high-affinity reagents (*k_off_* < 10^−4^ *s*^−1^) in 2D formats, where low CM densities result in equilibration times exceeding an hour **(Figure 1D)**. However, increasing [*C*] to 10 μM eliminates this dependence, driving *t*_99_ to sub-second timescales across the entire dissociation rate spectrum (10^−1^ *to* 10^−6^ *s*^−1^). This indicates that the 3D framework not only facilitates a higher capture fraction but does so with significantly reduced incubation times, a critical parameter for rapid point-of-care (POC) applications. Overall, this analysis also indicates that while the typical intuition that CM concentration does not matter in a binding assay as long as analyte concentration is much lower breaks down for weak binders where higher CM concentration can provide both equilibrium and kinetics advantages.

Finally, we addressed the relationship between signal enhancement and the analytical floor. While increasing [*C*] lifts the absolute signal, prior work has established that at sub-*K_D_* concentrations, the limit of detection (LOD) and the log-linear detection range are primarily governed by the background signal^4^. In traditional materials, increasing CM density often induces a proportional increase in non-specific binding, potentially negating sensitivity gains^5^. Consequently, realizing the full potential of volumetric capture requires a matrix capable of decoupling CM loading from background binding. This necessity informed our choice of a PEG-based hydrogel framework, whose bio-inert, hydrated backbone is intrinsically suited to suppress non-specific adsorption. By minimizing the background while maximizing capture efficiency, this material selection theoretically satisfies the core requirements for highly sensitive detection of low abundance analytes.

While these kinetic and equilibrium advantages represent the theoretical upper limit, a limitation of this analysis is that it does rely on the assumption that the binding reaction is not diffusion-limited, allowing us to treat the immobilized probes as a homogeneous volumetric concentration available for interaction with the target in the bulk solution, establishing the theoretical upper limit of performance. In practice, the transition to a 3D architecture can introduce potential mass-transport resistance^29^. Because the hydrogel matrix inherently restricts target diffusion, maintaining an open-pore structure is essential to minimize the deviation from these theoretical limits. This requirement necessitates the engineering of a highly porous architecture where the hydrogel mesh size is significantly larger than the hydrodynamic radius of the target.

### 2.2. Rational Design and Physical Characterization of the Bio-gels

The performance of a 3D hydrogel-based biosensing scaffold is governed by the interplay between the functionalization density of immobilized CMs and the matrix permeability to target analytes. A high density of capture sites is essential to maximize analyte binding, particularly for low-affinity interactions and at trace analyte concentrations. While the 3D volume of a hydrogel offers a high theoretical capacity for CM immobilization, this often comes at the cost of reduced permeability due to increased crosslinking density or steric hindrance within the network^22^. Such constraints can lead to diffusion-limited kinetics, where analytes accumulate only at the surface, leaving the bulk of the sensor underutilized and limiting sensitivity. To address this, we developed a formulation strategy designed to enhance analyte permeability as functionalization density is increased.

Using biotin as a model CM, we characterized functional biotinylated gel (Bio-gel) formulations composed of a PEG-4aNB (20 kDa) backbone, a bifunctional SH-PEG-SH (5 kDa) crosslinker, and a thiolated SH-PEG-Biotin (1 kDa) capture moiety **(Figure 2A)**. By maintaining a strict stoichiometric balance during synthesis, we increased the concentration of SH-PEG-Biotin to elevate functional density, which inherently reduced the degree of crosslinking in the resulting network. All tested Bio-gel formulations are detailed in Supplementary Table S1. We observed that a 50% reduction in the degree of crosslinking significantly increased the swelling ratios of the 10 wt% Bio-gels (referred to as 10/50; 10 wt% PEG-4aNB, 50% crosslinking) by 64% and the 6 wt% Bio-gels (6/50) by 102% **(Figure 2B)**. We identified the 6/50 Bio-gel as the optimal formulation, yielding soft yet stable hydrogels with a 46-fold swelling ratio, comparable to previously reported high swelling PEG formulations^30,31^. This architecture achieved a remarkably high local biotin loading of 6 mM, successfully realizing the high capture density and open-pore structure required for effective volumetric biosensing. Consequently, a 1 μL hydrogel spot, the standard detection zone utilized for all subsequent assays, has a loading capacity of 6 nmol of CMs. To put this in perspective, a traditional high-binding 96-well plate is limited to a maximum surface loading of approximately 1 pmol per well. By transitioning to a 3D framework, our Bio-gel platform thus provides over a 6000-fold theoretical increase in the immobilized capture moieties per sensing site, a capacity enhancement that aligns with previous reports for high-performance 3D diagnostic scaffolds **(Figure 2C)**^19^. Notably, this 50% crosslinking level represented a critical stability threshold, as formulations with lower crosslinking densities lacked the mechanical stability required for assay processing.

**Figure 2.**
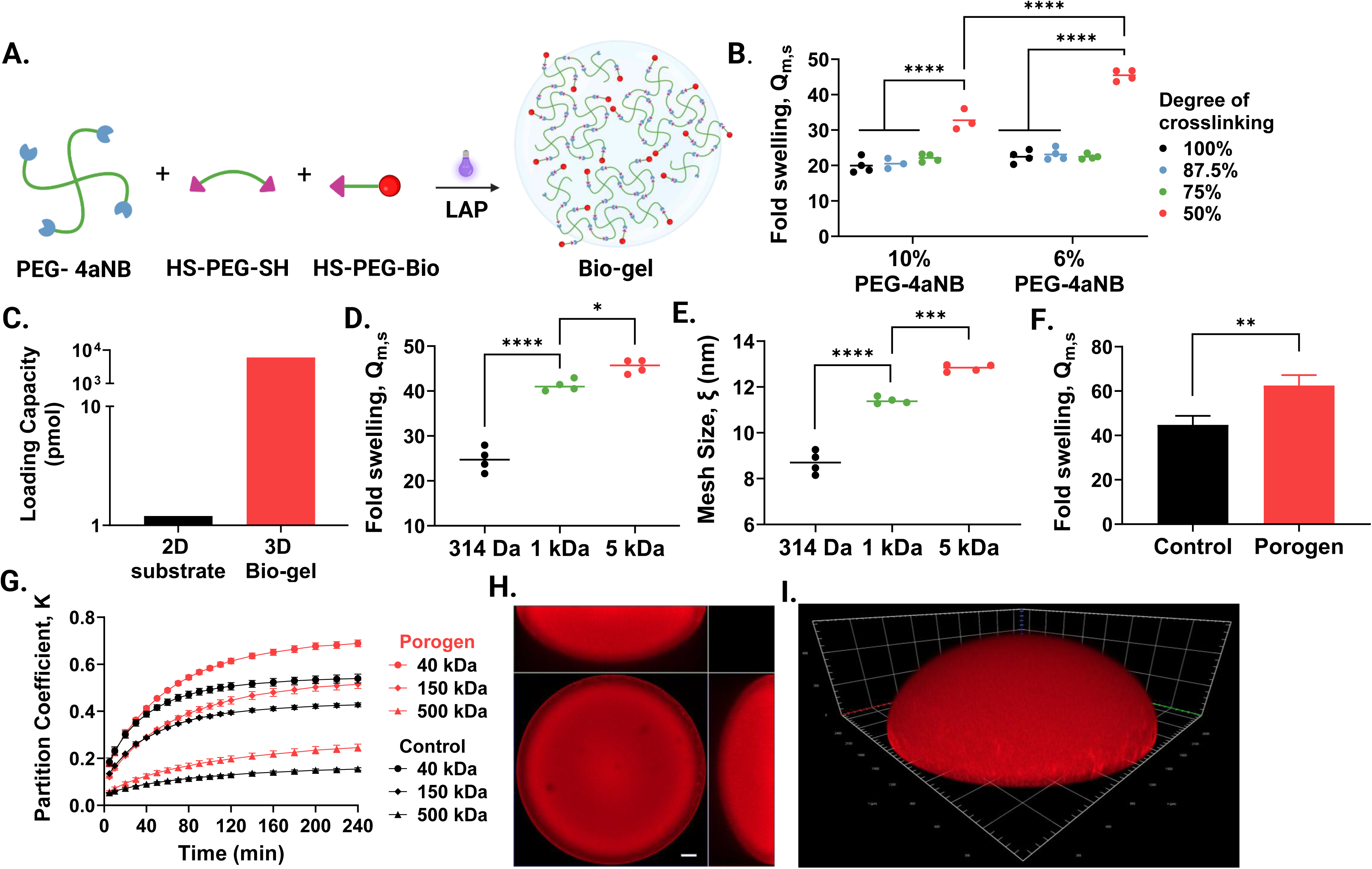
Rational design and physical characterization of functional PEG-based Bio-gels. **(A)** Schematic representation of the Bio-gel formulation, composed of a PEG-4aNB structural backbone, a bifunctional SH-PEG-SH crosslinker, and a thiolated SH-PEG-Biotin capture moiety. **(B)** Swelling ratios for 10 wt% and 6 wt% Bio-gel networks with varying degrees of crosslinking. **(C)** Theoretical loading capacity of the 3D platform compared to traditional 2D planar substrates. **(D)** Influence of crosslinker molecular weights (314 Da, 1 kDa, and 5 kDa PEG-dithiol) on Bio-gel swelling and **(E)** Estimated internal mesh size. **(F)** Swelling ratios of Bio-gels formulated with and without a non-reactive PEG 600 porogen (20% v/v). **(G)** Size-selective transport kinetics and partition coefficients of the 6/50 Bio-gel with and without porogen for FITC-Dextran of varying molecular weights. **(H)** Maximum intensity orthogonal projections of an optimized 1 μL Bio-gel detection spot tagged with fluorescently labelled streptavidin. **(I)** Confocal imaging of the spatial distribution of fluorescently tagged capture moieties within the hydrogel volume (Scale bar = 200 μm). All data are presented as the mean ± standard deviation (SD) for n = 3 to 5 experimental replicates. Statistical significance is denoted as *p ≤ 0.05, **p ≤ 0.01, ***p ≤ 0.001, and ****p ≤ 0.0001.

Using the 6/50 formulation as a baseline, we investigated the influence of crosslinker length on the internal mesh size (*ξ*). Increasing the crosslinker length significantly enhanced hydrogel swelling, highlighting its role as a critical parameter for optimizing 3D sensing performance **(Figure 2D)**^22^. Using modified Flory-Rehner equations, the mesh sizes were estimated at 8.7 nm, 11.4 nm, and 12.8 nm for PEG-dithiol crosslinkers with molecular weights of 314 Da, 1 kDa, and 5 kDa, respectively **(Figure 2E)**^32^. These physical dimensions are particularly significant when compared to the hydrodynamic radius (*R*_ℎ_) of IgG antibodies, a common diagnostic biomarker, which is approximately 5.5 nm (hydrodynamic diameter of ∼11 nm)^33^. While the 300 Da crosslinker results in a mesh (8.7 nm) that could sterically restrict the entry of IgG, the 5 kDa crosslinker provides a mesh size that significantly exceeds the analyte diameter, theoretically enabling enhanced volumetric diffusion and validating its use in the optimized formulation.

We further evaluated the addition of a porogen as a strategy to enhance matrix permeability. By incorporating PEG 600 (20% v/v) as a non-reactive porogen that is removed post-gelation, we aimed to introduce larger structural voids for improved analyte diffusion^18^ and observed a 50% increase in Bio-gel swelling **(Figure 2F)**. Transport kinetics for formulations with and without porogen were analyzed by monitoring the infiltration of FITC-Dextran with molecular weights of 40, 150, and 500 kDa (corresponding to *R*_ℎ_ of 4.3 nm, 7.9 nm, and 13.8 nm, respectively) into a 1 μL Bio-gel spot^34^. Corroborating with the mesh size estimate, our results demonstrated that while the 6/50 Bio-gel remains highly permeable to 40 and 150 kDa dextran – modelling a broad range of analytes including cytokines, growth factors, proteases and antibodies – it exhibits a significant reduction in the partition coefficient for 500 kDa species **(Figure 2G and Figure S1)**. This indicates that the engineered mesoporous framework of the hydrogel provides a size-selective sieving effect, potentially enhancing the signal-to-noise ratio by facilitating target analyte penetration while physically excluding larger, non-specific interferents often present in complex clinical matrices (e.g. red blood cells in blood). While the addition of PEG 600 improved dextran partition coefficients (27% for 40 kDa and 21% for 150 kDa), it ultimately compromised the long-term mechanical stability of the gels. Consequently, here, the porogen-free 6/50 formulation (referred to as Bio-gel henceforth) was selected for all subsequent experiments as it offered the optimum balance of functionality and structural stability. Diffusion kinetics studies confirmed that the partition coefficients for the 40 and 150 kDa species plateaued after 2 hours of incubation, informing the selection of a standardized 2-hour incubation period for all subsequent binding assays to ensure the system reached a transport equilibrium.

Finally, we characterized the spatial distribution of CMs within this optimized 1 μL Bio-gel detection spot. To visualize the biotin moieties, SH-PEG-Biotin was pre-conjugated with Strep-AF647 prior to photo-crosslinking. Confocal imaging revealed a roughly hemispherical geometry with a mean equilibrium diameter of approximately 2.5 mm and a height of 800 μm making them easily compatible with several high throughput microwell formats **(Figure 2H and Figure S2)**.

Crucially, the biotin molecules were found to be uniformly distributed throughout the entire volume of the hydrogel ensuring complete utilization of the detection spot for analyte capture **(Figure 2I)**.

### 2.3. Sensitivity Enhancement in 3D Fluorescence-Based Immunoassays

Following the physical optimization of the Bio-gel matrix, we evaluated its performance as a signal-amplifying scaffold using the high-affinity biotin-streptavidin interaction as a robust and quantifiable model system. To verify that the photo-crosslinking process did not compromise binding activity of the CM, Bio-gels were incubated with fluorescently tagged streptavidin to assess binding. After washing to remove unbound analyte, we observed a high, uniform fluorescence intensity throughout the gel matrix, validating the bio-orthogonal nature of the gelation chemistry and confirming that the crosslinking process did not denature the capture protein or obstruct its active binding sites **(Figure 3A)**. Furthermore, non-functionalized (NF-gel) controls displayed signal intensities indistinguishable from the background, even without the requirement for an additional blocking step, which is typically required for many 2D substrate-based immunoassays. This highlighted the inherent bio-inertness of the PEG framework and ensured that signal generation is strictly dependent on specific molecular recognition.

**Figure 3.**
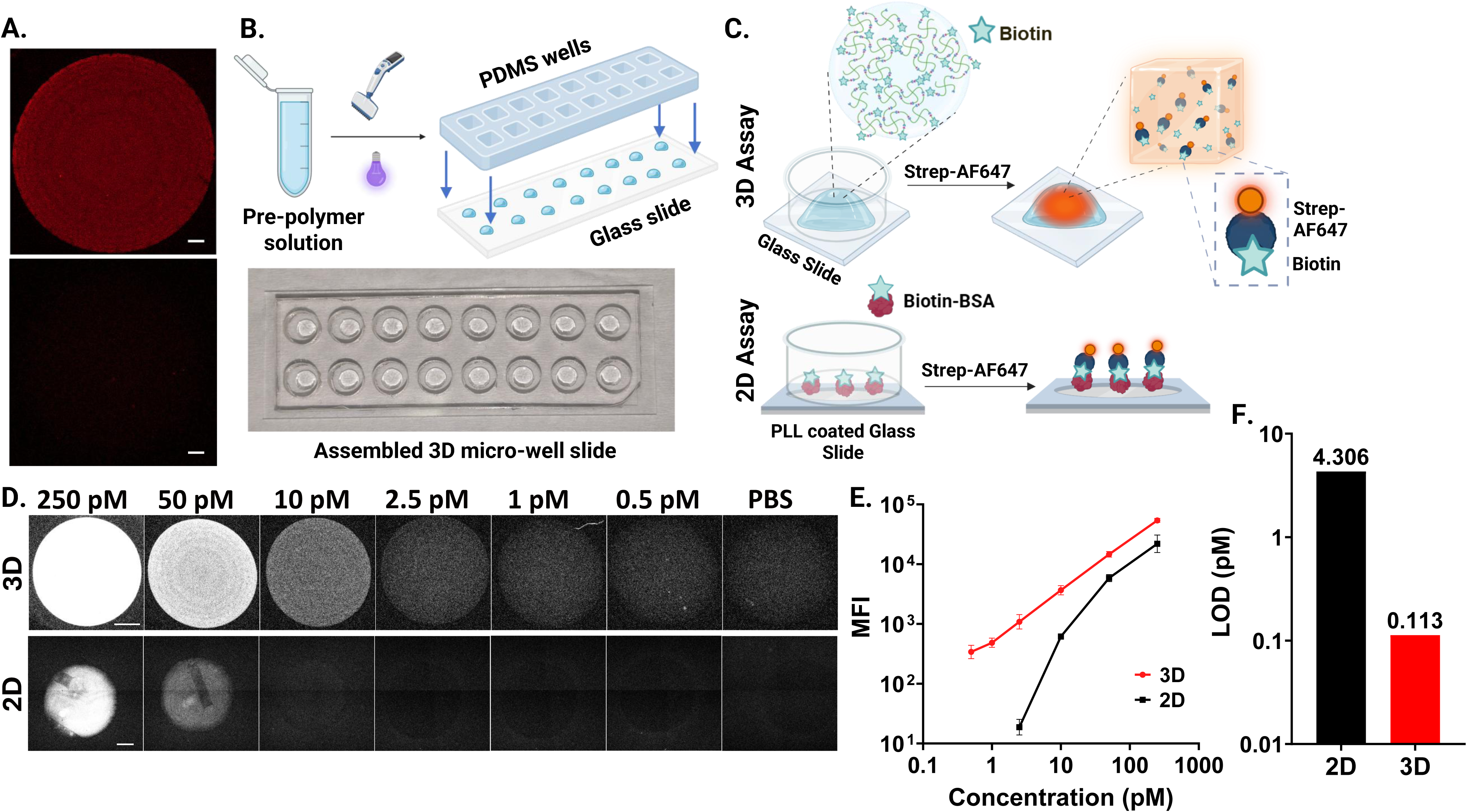
Sensitivity enhancement of fluorescence-based detection assays using 3D micro-well slide in a biotin-streptavidin model. **(A)** Fluorescence images of biotin functionalized hydrogel (Bio-gel) and non-functionalized (NF-gel) control following incubation with fluorescently tagged streptavidin (Scale bars = 200 μm). **(B)** Schematic representation of the assembly process for the 3D microwell slide and image of the assembled platform. **(C)** Schematic illustration of experimental workflows for the 3D and 2D biotin-streptavidin binding assays for fluorescence-based detection. **(D)** Fluorescence images of the 3D and 2D detection spots following incubation with serial dilutions of Strep-AF647 (Scale bars = 500 μm). **(E)** Serial dilution curves plotting mean fluorescence intensity (MFI) at the detection spot as a function of Strep-AF647 concentration for the 2D and 3D platforms. **(F)** Limits of detection for the 2D and 3D fluorescence-based detection assays. All data are presented as the mean ± SD for n = 3 experimental replicates.

To quantify the sensitivity enhancement provided by the hydrogel for streptavidin binding, we performed a comparison between the 3D Bio-gel and a conventional 2D substrate **(Figure 3B, C)**. The 3D platform consisted of 1 μL Bio-gel spots as volumetric detection zones arrayed onto a glass slide. For the 2D platform, a high biotin loading was achieved by immobilizing Biotin-BSA on poly-L-lysine (PLL) coated glass slides^35^. Unlike Bio-gel, the PLL surface was prone to non-specific adsorption and had to be blocked with BSA before the assay was performed. In both cases individual detection spots were physically isolated using reversibly sealed, leakproof PDMS reservoirs enabling multiple samples to be run on the same slide. To ensure a valid comparison, the 2D detection zone was standardized to a 2.5 mm diameter using PDMS functionalization templates to match the equilibrium diameter of Bio-gel spots.

Following incubation with serial dilutions of Strep-AF647, imaging of the detection spots revealed a stark divergence in detection capability between the 2D and 3D assay platforms **(Figure 3D)**. The 2D assay exhibited a rapid signal decrease with decreasing streptavidin concentration, reaching its analytical floor at concentrations below 10 pM. However, the 3D Bio-gel spots remained distinguishable from the baseline (PBS) even at sub-picomolar concentrations, maintaining a quantifiable response across a dynamic range spanning four orders of magnitude **(Figure 3E)**. Using the commonly three-sigma method^36^, we calculated the LOD for the Bio-gel to be 113 fM, representing a 38-fold improvement in sensitivity over the 2D control (LOD = 4.3 pM) **(Figure 3F)**. This significant sensitivity gain of the 3D platform is attributed to the synergistic effect of increased loading capacity and accessibility to the binding sites, allowing the concentration of a larger fraction of analyte within the detection zone and effectively amplifying the fluorescent signal relative to the background. Furthermore, the porous, hydrated environment of the Bio-gel facilitates a solution-like state for the immobilized biotin molecules. This ensures high steric accessibility and prevents the protein crowding or surface-induced denaturation frequently observed on rigid 2D substrates. These results highlight the potential of our functional PEG hydrogels as powerful signal-amplifying platforms capable of extending the detection limits of traditional binding assays.

### 2.4. Sensitivity Enhancement in Silver Metallization-based 3D Optical and Electronic Assays

While fluorescence-based assays offer high sensitivity, the requirement for complex optical instrumentation restricts their utility in resource-limited point-of-care (POC) settings. To overcome this, we have previously established an alternate sensing strategy using enzymatic silver metallization, which converts molecular recognition events into optically dense and conductive, dry-stable silver deposits and used it in diagnostics for Tuberculosis^37,38^ and Neglected Tropical Diseases^39^. This approach allows quantification using both using a cell phone camera and direct electronic sensing on nanostructured 2D electrodes, using a portable electronic reader or even a simple multimeter^35,40^. Although these systems successfully eliminate the need for complex optics, their analytical performance has been fundamentally bottlenecked by the finite analyte capture density of the 2D planar surfaces, restricting the total amount of silver that could be deposited.

Hypothesizing that our optimized Bio-gel serves as the ideal volumetric scaffold to overcome these 2D surface-bound constraints, we evaluated its performance in optical detection of silver metallization. Using the biotin-streptavidin model system, we employed horseradish peroxidase-conjugated streptavidin (Strep-HRP) as the analyte and catalytic reporter that drove reduction of silver ions into metallic silver deposits **(Figure 4A)**. Following the metallization reaction, the 1 μL Bio-gel spots generated significantly darker silver precipitates compared to 2D controls at equivalent concentrations **(Figure 4B)**. Crucially, the serial analyte dilution curve for 3D platform exhibited a significantly steeper slope compared to the planar control **(Figure 4C)**. This divergence indicates that for every incremental increase in analyte concentration, the volumetric matrix facilitates a larger accumulation of silver. Quantitative analysis revealed a LOD of 3 pM for the Bio-gel, representing a 23-fold improvement over the 2D assay **(Figure 4D)**. This enhancement is attributed to the porous structure of the Bio-gel, which allows silver aggregates to nucleate and grow three-dimensionally throughout the 800 μm scaffold height rather than being restricted to a thin surface, enabling superior optical discrimination.

**Figure 4.**
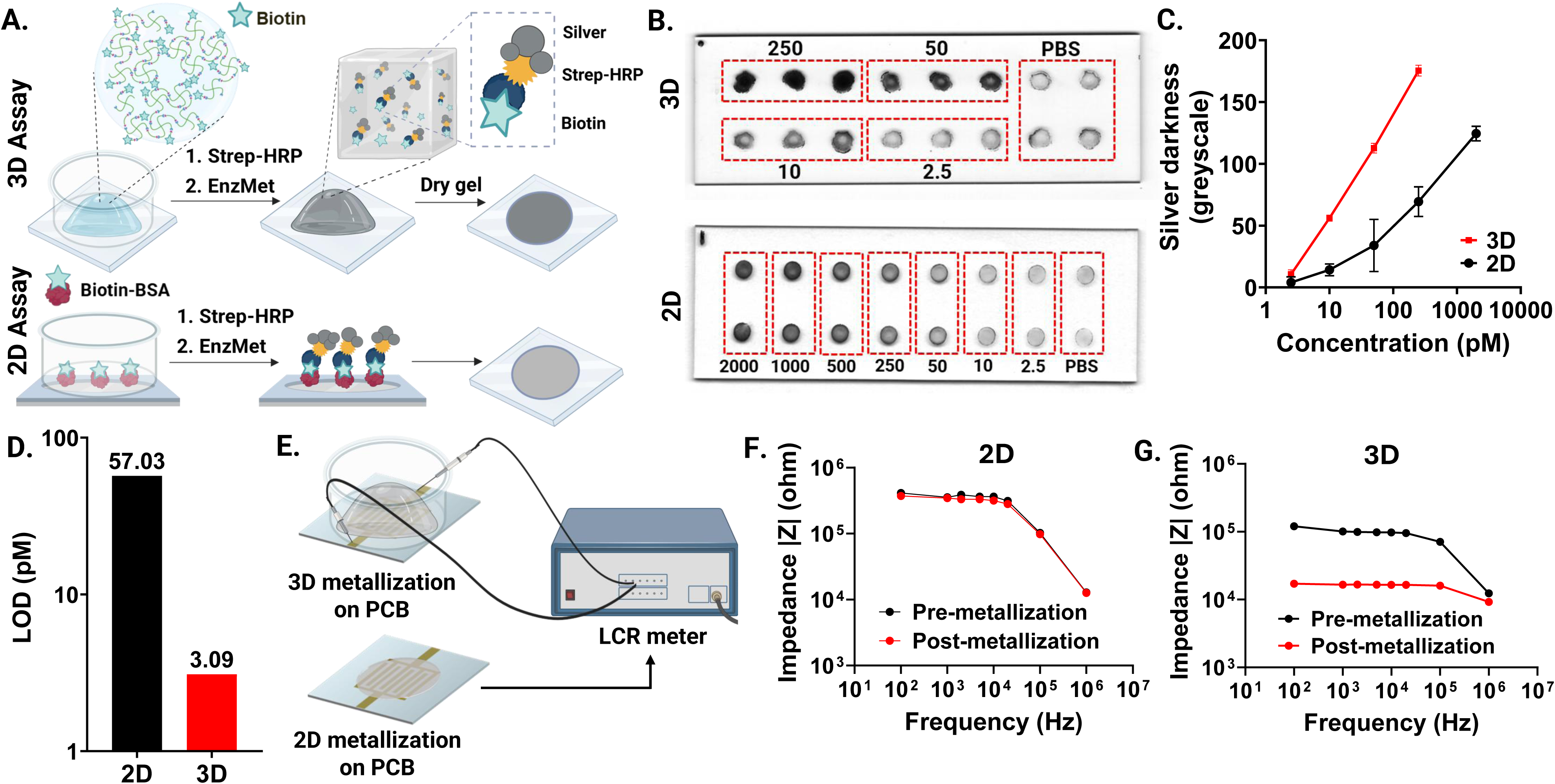
Sensitivity enhancement of enzymatic silver metallization-based optical and electronic detection assays using functionalized hydrogels in a biotin-streptavidin model. **(A)** Schematic illustration of the experimental workflows for the 3D and 2D biotin-streptavidin binding assays for optical detection. **(B)** Representative optical images of the silver deposits generated in the 3D and 2D detection spots following incubation with serial dilutions of Strep-HRP. **(C)** Serial dilution curves plotting optical darkness as a function of Strep-HRP concentration for the 2D and 3D platforms (n=3). **(D)** Limits of detection for the 2D and 3D optical detection assays. **(E)** Schematic illustration of the electronic sensing setup utilizing hydrogel modified (3D) and unmodified (2D) flexible interdigitated electrodes (FIDEs) on printed circuit boards (PCB). **(F)** Impedance response of 2D unmodified electrodes following incubation with 3 nM Strep-HRP and subsequent metallization. **(G)** Impedance response of 3D Bio-gel-modified electrodes following incubation with 3 nM Strep-HRP and subsequent metallization.

Building on these results, we next adapted the Bio-gel platform for electronic detection, a sensing modality that offers direct quantification without any intermediate optics and seamless integration with portable electronics **(Figure 4E)**^41^. Here, sensing relies on the probe-driven enzymatic reduction and deposition of conductive silver aggregates that bridge an open electrode circuit, leading to an analyte-dependent decrease in electrical impedance (|*Z*|). To evaluate performance on low-cost substrates, Bio-gel modified sensors were assembled on commercial flexible printed circuit boards (PCBs) with interdigitated electrodes (FIDEs) featuring a macroscopic electrode spacing of 0.25mm. While silver metallization was observed on both platforms, it failed to alter the impedance on unmodified 2D electrodes **(Figure 4F, Figure S3)**. On the other hand, Bio-gel-modified sensors registered a significant 10-fold reduction in |*Z*| **(Figure 4G)**. This confirms the inability of sparse surface-bound silver deposits to bridge wide electrode gaps and overcomes a critical prior limitation in this sensing chemistry that, in prior work, typically necessitates expensive, cleanroom-fabricated electrodes with narrow spacings (10µm)^42^ and the use of other enhancing catalytic agents such as gold nanoparticles^39^. The strong signal generated within the Bio-gel is driven by the formation of a dense, interconnected 3D network of silver aggregates throughout the scaffold. This volumetric percolation effectively shorts the electronic circuit formed across the 0.25mm gap, demonstrating that the 3D architecture eliminates the requirement for costly microfabrication for this detection method. While this initial demonstration serves as a proof-of-concept requiring further optimization for full electronic quantification, it represents, to the best of our knowledge, the first instance in which hydrogel-mediated silver metallization has been successfully translated into a quantifiable electronic measurement. By enabling high-sensitivity electronic sensing on inexpensive, mass-produceable substrates, this approach provides a scalable pathway for the development of sophisticated, digitally integrated point-of-care diagnostics.

### 2.5. Validation in Clinical Sample Matrices: Detection and Quantification of COVID-19 Antigen Specific Antibodies in Patient Serum

Having demonstrated significant sensitivity enhancements in model assays, we next validated the platform performance in a more directly clinically relevant application. To test the utility of our functional PEG hydrogels for disease diagnostics and monitoring, we selected the serological diagnosis of COVID-19 infection as the validation model, specifically targeting human IgG and IgA antibodies against the SARS-CoV-2 Nucleocapsid (NC) and Spike proteins (SP) as target biomarkers. A glass bottom 96-well plate, commonly used to perform fluorescence-based assays, was easily converted into a hydrogel enhanced bio-assay platform for nucleocapsid-specific IgG detection by simply pipetting the NC-gel formulation into each well followed by photo-crosslinking **(Figure 5A)**. The gel exhibited robust adhesion to the well surface without the need for any surface modification steps highlighting the ease of integration of our hydrogel formulation with commonly used high throughput assay formats.

**Figure 5.**
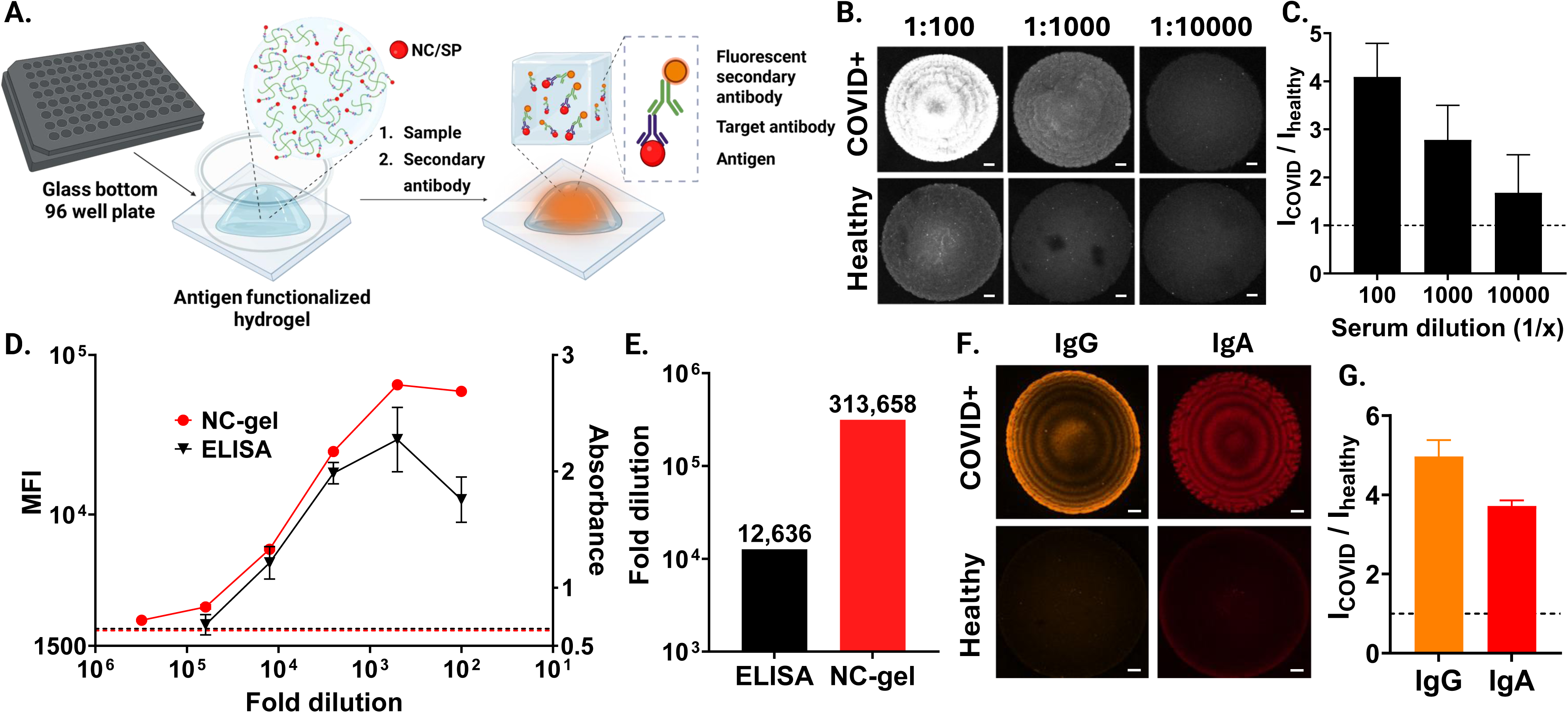
Detection and quantification of COVID-19 antigen-specific antibodies in clinical samples. **(A)** Schematic representation of hydrogel-enhanced immunoassay for nucleocapsid (NC) or spike protein (SP)-specific IgG detection in a 96-well plate format. **(B)** Fluorescence images of nucleocapsid-functionalized hydrogels (NC-gels) following incubation with serial dilutions of pooled COVID-19 positive patient serum and pre-pandemic healthy control serum (Scale bars = 200 μm). **(C)** Plot of the ratio of fluorescence intensity (COVID-19 positive to healthy control) as a function of serum dilution for the NC-gels. **(D)** Serial dilution curves comparing the signal of NC-gel assay to a standard ELISA across dilutions of COVID-19 positive patient serum. **(E)** Limits of detection for the NC-gel assay and ELISA. **(F)** Fluorescence images of spike protein-functionalized hydrogels (SP-gels) following incubation with patient serum and a cocktail of distinct secondary antibodies against IgG and IgA (Scale bars = 200 μm). **(G)** Resolved fluorescence signal intensities for the multiplexed detection of IgG and IgA within a single 1 μL detection spot. All data are presented as the mean ± SD for n = 3 experimental replicates.

We first evaluated the ability of the NC-gels to discriminate between diseased and healthy samples in a complex biological matrix. NC-gels were incubated with serial dilutions of pooled COVID-19 positive patient serum and pre-pandemic healthy controls, followed by probing with fluorescently tagged anti-human IgG. Imaging revealed that the platform maintained its high specificity with the gels clearly differentiating patient samples from healthy controls at serum dilutions as low as 1:10,000 **(Figure 5B and C)**. This is consistent with the bio-inertness of the PEG scaffold that reduces non-specific binding and trapping of matrix components or probes within the detection spot. The low yet discernable fluorescence observed for the healthy serum can be attributed pre-existing cross-reactive antibody responses to endemic human coronaviruses which has been described in previous work on antibody profiling in COVID-19^43^.

Next, we quantified the sensitivity enhancement provided by the NC-gels relative to a standard optical ELISA, a commonly used assay in clinical serology – replicating the 3D vs 2D comparisons from our model assay – with a more complex sample type and assay chemistry **(Figure 5D)**. We observed that on incubation with serial dilutions of pooled COVID-19 patient samples, the ELISA signal decayed to baseline levels at dilutions below 1:12,500. In contrast, the NC-gels offered a wider dynamic range, generating signal higher than the buffer-only baseline at dilutions as low as 1:312,500, achieving a 25-fold improvement in LOD over the clinical gold standard **(Figure 5E)**. Clinically, such ultrasensitive detection has implications for identifying seroconversion in early-stage disease or monitoring low-abundance biomarkers, such as cytokines, where concentrations can fall below the LOD of conventional 2D assays. It may also enable the use of more dilute serum samples, which may reduce matrix effects and may conserve valuable clinical specimens. Furthermore, the NC-gels exhibited a significantly reduced “hook effect” (prozone phenomenon) and generated higher signal-to-noise ratios at lower serum dilutions compared to ELISA **(Figure 5C)**. This is due to the expanded binding capacity of the 3D matrix, which prevents early saturation of available capture sites which is a common limitation on 2D surfaces. While the exact concentration of polyclonal NC-specific IgG in the pooled serum is variable, antigen-specific IgG is estimated to comprise up to 1% of total serum IgG (∼10 mg mL^-1^) in high-titer samples^44^. At the limiting dilution of 1:313,000, this corresponds to a NC-specific IgG concentration of approximately 2 pM. This aligns well with the LODs established in our biotin-streptavidin model assays, confirming that the high sensitivity of the platform is preserved even in complex matrices such as clinical serum samples.

Finally, we utilized the expanded volumetric capacity of the Bio-gel to achieve simultaneous multi-analyte profiling in a single well, overcoming a limitation inherent to traditional ELISAs. To evaluate the ability of the platform to resolve diverse antibody isotypes, Spike protein-functionalized hydrogels were incubated with SARS-CoV-2 patient serum (1:100 diluted) and probed with a cocktail of secondary antibodies targeting IgG and IgA labeled with spectrally distinct fluorophores. Imaging resolved clear, independent signals for both isotypes within the same 1 μL detection spot **(Figure 5F and G)**. This simultaneous capture of IgG and IgA demonstrates that the mesoporous 3D matrix provides sufficient physical space to mitigate the steric crowding, site saturation and competitive binding typically observed on 2D surfaces and accommodates simultaneous binding events throughout the depth of the scaffold. By enabling high-content multiplexed serological profiling in a single well, this volumetric approach reduces sample volume requirements and workflow complexity, establishing our functional hydrogel as a robust formulation for complex, multiplexed diagnostic applications.

### 2.6. Performance Enhancement in Hydrogel-Integrated LFAs

Despite their ubiquity in point-of-care diagnostics, LFAs are frequently hindered by poor analytical sensitivity at low analyte concentrations, a lack of quantitative reliability and multiplexability^45^. We evaluated the potential of our functionalized hydrogel to enhance the performance of these simple, rapid, and inexpensive LFAs by integrating the photo-crosslinkable precursor directly into the nitrocellulose membrane commonly used for LFAs. Upon UV exposure, the hydrogel network polymerizes within the porous architecture of the NC substrate, creating a mechanically intermeshed composite detection zone^26^. To benchmark this platform, we utilized a Protein A/G capture assay for the detection of fluorescently tagged IgG as a model system reflecting the standard control line in commercial LFAs **(Figure 6A)**.

**Figure 6.**
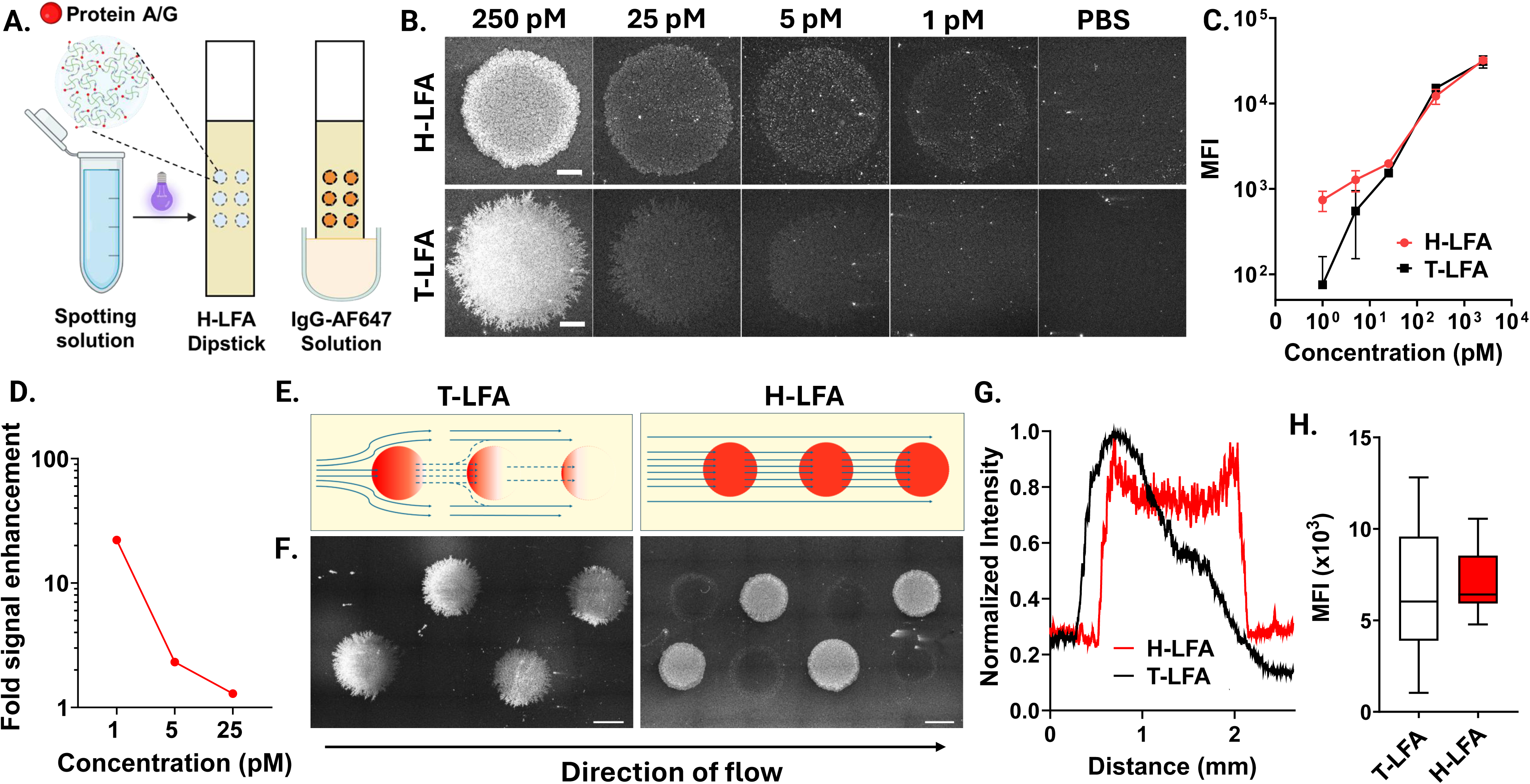
Performance enhancement and spatial multiplexing in hydrogel-integrated lateral flow assays (H-LFAs). **(A)** Schematic of the spotting process for H-LFA dipsticks with protein A/G functionalized hydrogel (AG-gel) and the binding assay for capturing fluorescently tagged IgG. **(B)** Fluorescence images of H-LFA and traditional LFA (T-LFA) detection spots following the application of serial dilutions of IgG-AF647 (Scale bars = 500 μm). **(C)** Serial dilution curves plotting mean fluorescence intensity (MFI) at the detection spot as a function of IgG-AF647 concentration for the H-LFA and T-LFA platforms (n=3 per condition). **(D)** Calculated fold-enhancement of the fluorescence signal in the H-LFA relative to the T-LFA. **(E)** Schematic illustrating the downstream shadow effect in a T-LFA with multiple detection spots and its resolution in the H-LFA. **(F)** Fluorescence images of the 2 x 4 T-LFA and H-LFA arrays consisting of alternating Protein A/G and BSA spots following IgG-AF647 capture (Scale bars = 1 mm). **(G)** Normalized fluorescence intensity profiles of an individual T-LFA and H-LFA capture spot in the direction of fluid flow. **(H)** Comparing variation in fluorescence intensity across the functionalized spots within the 2 x 4 T-LFA (n=10) and H-LFA (n=12) arrays.

A comparison between Protein A/G functionalized hydrogel (AG-gel) spotted LFAs (H-LFA) and traditional Protein A/G LFAs (T-LFA) across a serial dilution of fluorescently tagged IgG (1 pM to 2500 pM) revealed that the H-LFA produced significantly brighter and more well-delineated binding spots, particularly at trace analyte concentrations **(Figure 6B and C)**. The H-LFA achieved a 20-fold signal enhancement over T-LFAs at the lowest tested concentration of 1 pM **(Figure 6D)** while both platforms converge at higher analyte concentrations. This superior sensitivity can be attributed to the local alteration of fluid dynamics within the composite zone. By intermeshing with the membrane fibers, the hydrogel creates a more tortuous flow path and a localized reduction in effective membrane pore size, which slows the flow velocity, increases analyte residence time and improves analyte binding^25,26^. Furthermore, the use of flexible PEG linkers for Protein A/G immobilization ensures higher steric freedom and better orientation of capture moieties compared to the random, rigid adsorption on nitrocellulose surfaces^46^. Morphologically, the H-LFA produced perfectly circular detection spots **(Figure 6B)**. These facilitate precise quantification through standardized circular regions of interest (ROI), enabling easier automation, whereas T-LFA spots exhibited a heterogeneous, dendritic appearance. This advantage stems from the high hydrophilicity of the PEG hydrogel, which promotes uniform wetting and controlled wicking during the spotting process.

An additional unique advantage of this 3D integration was observed in the context of spatial multiplexing on LFAs. The development of lateral flow microarrays has emerged as a critical strategy for high-content diagnostics, enabling the simultaneous detection of multiple pathogens or biomarkers from a single patient sample^47,48^. A commonly observed yet rarely reported phenomenon in these microarrays is the depletion of binding signals in the direction of flow resulting in signal heterogeneity and unreliable quantification across the array **(Figure 6E)**^49–52^. We observed this in our 2 x 4 array of alternating Protein A/G and non-functionalized spots, where T-LFA arrays exhibited a “shadow effect,” leaving downstream locations with weakened signals **(Figure 6F).** In contrast, the H-LFAs demonstrated uniform signal intensity across all functionalized zones, regardless of their position in the flow path. Furthermore, each spot in the T-LFA array depicted a front-loaded “crescent effect,” where a distinct signal gradient was observed within the spot **(Figure 6G)**. This gradient was markedly absent in the H-LFA where the signal was uniformly distributed throughout the spot. Both effects occur in T-LFAs because the analyte accumulation at the front end of the capture zone causes it to act as impermeable pillar splitting the capillary flow. This causes analyte-laden fluid to diverge around the spot rather than passing completely through it and consequently, downstream capture spots display lower signals not necessarily because the total analyte is depleted, but because of this local flow obstruction. We hypothesize that the hydrogel modification, owing to its hydrophilicity, overcomes this effect by re-engineering the local fluid field and ensuring an even distribution of the analyte throughout the detection spot. Statistical analysis confirmed this uniformity, with the H-LFA showing a significantly lower standard deviation in fluorescence intensity across the array than the T-LFA **(Figure 6H)**.

While previous research has utilized hydrogels as structural tools for porosity tuning on paper-based substrates, this study represents, to the best of our knowledge, the first report of functionalized hydrogel scaffolds being used for the immobilization of CMs within nitrocellulose and to overcome the shadow/crescent effects. Ultimately, by overcoming the physical limitations of shadowing while increasing analyte capture signal, these intermeshed composite H-LFAs have the potential to enable the reliable and simultaneous measurement of multiple biomarkers from a single low-volume sample. This scalable technology provides a flexible solution for next-generation point-of-care diagnostics, where high-content profiling of antibody isotypes or Fc profiles^37,53,54^ enabling resolution of different disease states, current vs past infection, or co-infecting pathogens is required without sacrificing the simplicity of the standard lateral flow format which has been optimized for inexpensive mass manufacturing ($0.1-$1 per test) already. This can also be compared to the flow-through immunoassays or immunofiltration assays, which have been more recently called vertical flow assays as well, that have been developed over the last few decades for multiplexing of various analytes ^51,55–58^.

## 3. Conclusion

This work establishes a versatile PEG-based hydrogel platform designed to circumvent the fundamental physical and analytical constraints inherent to 2D substrate-based biosensing. By transitioning from a planar, surface-bound model to a volumetric 3D matrix, our functionalized hydrogels enable a 6000-fold increase in CM loading without compromising analyte permeability. By leveraging a porosity-tunable framework, the platform ensures that capture sites remain sterically accessible and uniformly distributed throughout the sensing volume. This architecture facilitates efficient analyte infiltration and volumetric utilization, effectively resolving the issues of steric crowding and site masking that frequently restrict the performance of capture moieties on rigid, 2D interfaces.

Central to this performance is the bio-orthogonal nature of the thiol-ene photo-click crosslinking strategy. The modularity of this specific reaction chemistry facilitates the covalent immobilization of virtually any thiolated biorecognition element. Given that thiolation is a ubiquitous and accessible modification for common capture probes—including antibodies, aptamers, and other nucleic acids—this platform provides a generalizable route for their seamless integration into the 3D matrix without compromising native conformation or obstructing active binding sites. Furthermore, the use of photo-initiated crosslinking provides precise temporal control over the polymerization process, including potential for photopatterning. Unlike conventional chemical gelation, which is often limited by narrow, time-sensitive processing windows, photo-initiation allows precursor solutions to be dispensed across high-density arrays or multi-well plates before undergoing concurrent gelation via a single UV exposure. This scalability is essential for high-throughput fabrication, where the detection zones can be further miniaturized through droplet printing or high-precision 3D bioprinting to enable high-density, truly multiplexed microarrays on both glass and paper-based substrates^59^.

The results presented here across multiple sensing modalities confirm the universal applicability of this signal-amplifying scaffold. The 3D matrix achieved a 38-fold improvement in fluorescence sensitivity, without the need for any blocking or related washing steps, and a 23-fold gain in silver metallization-based optical detection compared to conventional 2D baselines. While fluorescence microscopy was utilized here for quantification, the ongoing development of portable, cost-effective fluorescence readers provides a clear pathway to eliminate the need for expensive optics, ensuring these sensitivity gains remain accessible in decentralized settings for point-of-care use as well^60,61^. Furthermore, the integration with electronic sensing on inexpensive flexible electrodes demonstrated a fundamental shift in device physics, highlighting the first reported instance where the extended network of silver aggregates in a 3D matrix enabled readout on inexpensive, macroscopic flexible electrodes. This effectively eliminates the requirement for high-precision nanofabrication, paving the way for low-cost, digitally integrated diagnostics.

Clinical validation using COVID-19 serology bolsters the rigor of the platform, providing a 25-fold sensitivity improvement over the clinical gold-standard ELISA. The ability to decouple IgG and IgA responses within a single sensing volume demonstrates a powerful capacity for high-content profiling that conserves low volume clinical specimens. Similarly, the adaptation of this technology into LFAs highlights the first reported instance where a functionalized hydrogel is used as a matrix for immobilizing CMs within nitrocellulose. Furthermore, by re-engineering the local fluid dynamics to mitigate downstream signal depletion, the hydrogel-modified LFA architecture establishes the potential for reliable, quantitative multiplexing within standard lateral flow formats. Beyond traditional nitrocellulose, the mechanical intermeshing of the hydrogel network can enable the effective immobilization of proteins and nucleic acids on alternative, lower-cost paper substrates as well. This approach is particularly advantageous for materials such as cellulose filter papers, which are ubiquitous in paper microfluidics, typically lack the inherent binding capacity required for high-performance assays.

Despite these analytical enhancements, translating hydrated polymeric networks into point-of-care devices presents specific physical limitations that will need to overcome in future work before clinical translation can occur. Primarily, ambient storage of highly hydrated matrices is challenging; dehydration can induce structural collapse, while prolonged wet storage risks gradual polymer and capture molecule degradation. Further work is required to rigorously evaluate long-term wet storage stability and identify optimal storage conditions or stabilizing additives such as humectants and preservation buffers to maintain functional and structural integrity over time. Furthermore, while the current porosity successfully accommodates typical protein biomarkers, the diffusion of macro-scale targets (e.g., whole pathogens, intact cells, or large extracellular vesicles) into the dense 3D matrix remains sterically hindered. Extending this volumetric approach to cellular diagnostics^62,63^ will require engineering macro-porous networks. This could be achieved by utilizing Microporous Annealed Particle (MAP) scaffolds to create continuous interstitial voids, or through gas-foaming or cryogelation approaches for pore generation.

Ultimately, this platform establishes a universal methodology for the enhanced functional immobilization of CMs across a broad spectrum of diagnostic substrates. The modular nature of the hydrogel makes it inherently adaptable to a wide array of biomarkers beyond antiviral antibodies, including cytokines, proteases, and other small-molecule targets. By providing a standardized, ‘plug-and-play’ 3D matrix that integrates seamlessly into multi-well plates, glass slide-based microarrays, nitrocellulose membranes or electronic devices, this technology bridges the gap between laboratory-grade analytical performance and field-deployable low cost and high speed. Future work will focus on scaling this modular approach and translating these sensitivity gains to address critical needs in global health and exploring the development of stimuli-responsive hydrogel matrices. By eliminating the need for blocking or multiple washes and probe addition steps, such platforms could enable “add-and-read” diagnostic formats providing a truly POC solution for infectious disease monitoring in resource-limited settings.

## 4. Experimental Methods

### 4.1. Materials

4-arm PEG-norbornene (PEG-4aNB, 20 kDa) and PEG-dithiol (SH-PEG-SH, 5kDa) were obtained from JenKem Technology. Thiol-PEG-Biotin (SH-PEG-Bio, 1 kDa) and Thiol-PEG-Succinimidyl Carboxymethyl Ester (SH-PEG-SCM, 1 kDa) were obtained from BioPharma PEG. PEG-dithiol (314 Da and 1 kDa), PEG 600, FITC labelled dextran (4kDa, 150 kDa and 500 kDa) and bovine serum albumin (BSA) were obtained from Sigma Aldrich. Poly-L-lysine (PLL) coated slides, deionized (DI) water, phosphate buffer saline (PBS), HEPES (1M) buffer, biotinylated BSA (Biotin-BSA), horseradish peroxidase conjugated streptavidin (Strep-HRP), Alexa Fluor 647 conjugated streptavidin (Strep-AF647), polydimethylsiloxane (PDMS), TMB substrate and recombinant Protein A/G were obtained from Thermo Fisher Scientific. Nitrocellulose membrane (HF07502XSS) and cellulose absorbent pad were obtained from Millipore Sigma. Tween 20 was obtained from VWR. PDMS sheet was purchased from Greene Rubber Company. Lithium phenyl-2,4,6-trimethylbenzoylphosphinate (LAP) was obtained from Tocris Bioscience. SARS-CoV-2 nucleocapsid protein (NC, 40588-V07E) and spike protein (SP, 40604-V08B) were obtained from Sino Biological. EnzMet for General Research Applications was obtained from CedarLane. Mouse anti-human IgG Fc-HRP, goat anti-human IgG-PE, goat anti-human IgG-AF647 and goat anti-human IgA-AF647 were obtained from Southern Biotech. Flexible printed circuit boards with interdigitated electrodes (FIDE, SEN0004) were obtained from DigiKey.

### 4.2. Clinical Samples

COVID-19 positive and pre-pandemic healthy human serum samples were procured from Ray Biotech and Jackson Immuno, respectively. All clinical specimens were collected from donors who provided informed written consent prior to the sampling process (PROTOCOL NO: SOP-TF-PH-002 STERLING IRB ID: 8291-BZhang). Details are on file and available with them.

### 4.3. Synthesis and Bio-functionalization of PEG-based Hydrogels

Biotin-functionalized (Bio-gels) and non-functionalized (NF-gels) hydrogels were synthesized using 20 kDa PEG-4aNB as the structural backbone. The macromers were crosslinked with SH-PEG-SH of varying molecular weights (314 Da, 1kDa, or 5 kDa), with 1 kDa Da SH-PEG-Bio incorporated as the functional moiety. Pre-polymer solutions were prepared by dissolving PEG-4aNB (6 or 10 wt%) in PBS containing 10 mM HEPES and 1 mM LAP as the photoinitiator. The stoichiometric ratios of SH-PEG-SH and SH-PEG-Bio were adjusted to achieve crosslinking degrees of 50, 75, 87.5, or 100%, where the 100% formulation represented the NF-gel control. To enhance hydrogel porosity, 20% v/v PEG 600 was added to the Bio-gel formulations as a porogen. All formulations were crosslinked via UV exposure (365 nm, UWAVE, France) at 50 mW cm^-2^ power for 45 s.

To generate hydrogels functionalized with SARS-CoV-2 NC protein (NC-gels), SP protein (SP-gels), or Protein A/G (AG-gels), the respective CMs were first thiolated using a heterobifunctional PEG linker via NHS chemistry to enable covalent conjugation to the PEG-4aNB backbone. Briefly, 100 μL of the CM in PBS was incubated with 8 μL of 250 mM SH-PEG-SCM in DMSO for 2 hours. These thiol-labeled CMs (SH-PEG-CM) were then incorporated into formulations containing 3 or 6 wt% PEG-4aNB and 5 kDa SH-PEG-SH to produce functionalized hydrogels with a 50% degree of crosslinking. The crosslinking duration for the 3 wt% and 6 wt% formulations was 3 min and 45 s, respectively. via NHS chemistry to enable covalent conjugation to the PEG-4aNB backbone.

### 4.4. Hydrogel Swelling and Mesh Size Calculations

The swelling behavior and network structure of the various hydrogel formulations were quantified to assess the influence of polymer concentration, crosslinking agent and porogen addition on the resulting matrix porosity. For swelling characterization, 20 μL of the precursor solution was crosslinked, and the initial mass (Wi) was recorded immediately after preparation. The hydrogels were then incubated overnight in PBS at room temperature (RT) to reach equilibrium swelling (*n* = 5 per condition). Following incubation, excess surface water was removed, and the equilibrium swollen mass (Ws) was recorded. The samples were subsequently dehydrated at 65°C for 24 hours to remove all water content, and the final dry mass (Wd) was measured.

The mass swelling ratio or fold swelling in the equilibrium swollen state (*Q_m_*_,*s*_) and relaxed state (*Q_m_*_,*r*_) were calculated as Ws/Wd and Wi/Wd, respectively. These values were further used to calculate the volumetric swelling ratios in the equilibrium (*Q_v_*_,*s*_) and relaxed (*Q_v_*_,*r*_) states as follows:

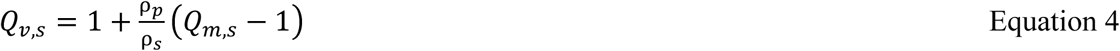

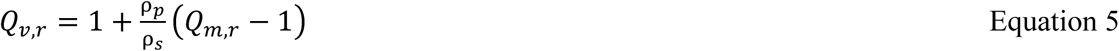

Here, *ρ_p_* is the density of the polymer (1.12 g cm^-3^ for PEG) and *ρ_s_* is the density of the solvent (1.0 g cm^-3^ for PBS). The polymer volume fractions in the swollen state (*v*_2,*s*_) and relaxed state (*v*_2,*r*_) were then determined as reciprocals of *Q_v_*_,*s*_ and *Q_v_*_,*r*_ respectively.

The molecular weight between crosslinks 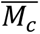 was calculated using the Peppas-Merrill model, a modification to the Flory-Rehner equation^32^

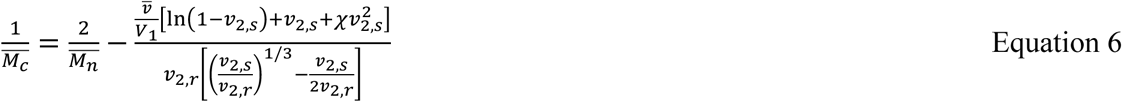

where 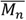 is the number-average molecular weight of the uncrosslinked hydrogel components, 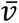 is the specific volume of the polymer (*ρ_s_*/*ρ_p_*), *V*_1_ is the molar volume of water (18 cm^3^ mol^-1^), and *χ* is the Flory-Huggins polymer-solvent interaction parameter (0.426 for PEG-water).

Finally, the mesh size (*ξ*) was determined as described by Canal and Peppas^64^

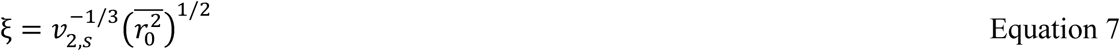

where the unperturbed end-to-end distance 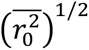 was determined based on the PEG backbone properties as follows:

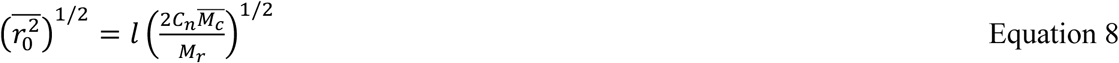

Here, *l* is the average bond length of the PEG backbone (0.146 nm), *C_n_* is the Flory characteristic ratio (4.0 for PEG), and *M_r_* is the molecular weight of the PEG repeat unit (44 g mol^-1^)^65^.

### 4.5. Characterizing of FITC-dextran Diffusion into Hydrogel Matrix

To evaluate the molecular transport properties and the effect of porogen-induced porosity, diffusion experiments were performed using FITC-labeled dextrans of varying molecular weights (40, 150, and 500 kDa). Bio-gel droplets (1 μL), with and without the addition of PEG 600 porogen, were dispensed into an ibidi 18-well μ-slide and crosslinked. The gels were incubated in PBS overnight to achieve equilibrium swelling and to ensure the complete removal of unbound porogen. 150 μL of a 50 μg mL^-1^ FITC-dextran solution in PBS was added to each well (*n* = 3 per condition) and time-lapse imaging was initiated immediately using a fluorescence microscope (LSM 900, Zeiss, Germany). Images were acquired every 10 minutes at a fixed *z*-height of 100 μm from the base of the gel to monitor the infiltration of the fluorescent probes into the hydrogel matrix. At each time point, the partition coefficient (*K*) was determined using the following:

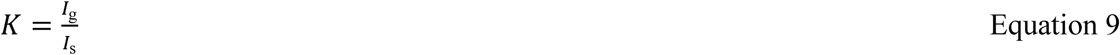

Here, *I_g_* represents the mean fluorescence intensity within the hydrogel matrix and *I_s_* represents the mean fluorescence intensity of the surrounding FITC-dextran solution quantified using ImageJ.

### 4.6. Characterizing Distribution of CMs within Hydrogel Detection Spot

To verify homogeneous distribution of the CMs within the functionalized Bio-gel and characterize the physical dimensions of the hydrogel detection spot, a fluorescent labeling strategy was employed. SH-PEG-Bio was pre-conjugated with Strep-AF647 prior to its incorporation into the polymer network. Briefly, 1 mg of SH-PEG-Bio was incubated with 33 μL of Strep-AF647 (2 mg mL^-1^) for 1 hour at RT before integrating into the Bio-gel formulation. 1 μL droplets of the labeled formulation were dispensed onto an ibidi 18-well μ-slide (*n* = 4), crosslinked, and equilibrated in PBS overnight. After washing to remove unbound fluorophores, confocal microscopy (Zeiss 900A) was used to acquire a z-stack of the hydrogel spot. Images were captured at 20 μm intervals along the *z*-axis to obtain a 3D reconstruction of the distribution of biotin within the gel. The equilibrium diameter and height of the detection spot was further calculated using ImageJ, providing the standardized geometric footprint used for all subsequent experiments.

### 4.7. PDMS Component Fabrication

PDMS membranes were fabricated to define the sensing geometries and sample volumes for the detection assays. A thin PDMS film (0.25 mm thickness) and a cast PDMS sheet (2.5 mm thickness) were patterned using a CO2 laser (PLS6.150D, Universal Laser Systems, USA) to create arrays of 2.5 mm and 6 mm diameter wells, respectively. These components served as the functionalization template (thin layer) and the sample reservoir (thick layer) respectively in subsequent microwell assays. The parts were soaked in 5% Alconox aqueous solution and bath-sonicated for 10 min to remove any carbonaceous residue, rinsed thoroughly with DI water and air-dried. Prior to assembly, adhesive tape was used to remove any residual microparticles or debris from the surface to ensure a leak-proof seal.

### 4.8. 2D Fluorescence Assay

A 2D biotin-streptavidin model binding assay was conducted to provide a comparative baseline for the 3D hydrogel sensing platform. To prepare the assay substrate, the thin PDMS functionalization template was reversibly sealed onto a PLL-coated glass slide. To functionalize the exposed regions with biotin, 1 mg mL^-1^ of biotin-BSA in PBS was added to each well and incubated for 1 hr at RT in a humidified chamber. Following incubation, the wells were blocked with 1% BSA in 0.1% Tween 20 in PBS (0.1% PBST) for 30 min and washed twice with 0.1% PBST and once with PBS. All washing steps were performed by placing the glass slide in a petri dish filled with washing buffer on a plate shaker for 10 min at 100 rpm.

The thicker PDMS reservoir layer was then aligned and reversibly sealed over the existing template, creating leak-proof wells to hold the analyte solutions. Serial dilutions of Strep-AF647, ranging from 0.5 pM to 250 pM were prepared in 1 mg mL^-1^ BSA in 0.05% PBST. A 100 μL volume of each dilution was added to the reservoir wells (*n* = 3 per condition) and incubated for 2 h on a plate shaker at 60 rpm. Following incubation, the slides were washed as detailed above, rinsed with DI water, and air-dried. The resulting fluorescence intensity of the bound Strep-AF647 was acquired using fluorescence microscopy and quantified via ImageJ.

### 4.9. 3D Fluorescence Assay

To evaluate the sensitivity enhancement provided by the 3D hydrogel matrix, the biotin-streptavidin binding assay was performed Bio-gel enhanced assay substrates. First, the thick PDMS reservoir layer was reversibly sealed onto a standard microscope glass slide. A 1 μL droplet of the Bio-gel precursor solution was pipetted into the center of each well and photo-crosslinked to immobilize the biotinylated capture network. The resulting 3D detection spots were incubated in PBS overnight to reach equilibrium swelling. No additional blocking step was required.

To assess analyte capture, detection spots were incubated with serial dilutions of Strep-AF647 (*n* = 3 per condition) and washed according to the protocol detailed previously. Post washing, the wells were filled with fresh PBS, and the captured streptavidin was quantified via fluorescence microscopy. For each detection spot, a z-stack was acquired consisting of 12 images at 80 μm intervals from the base of the gel to a total height of 880 μm. Maximum intensity projections (MIPs) were generated from these stacks to represent the total fluorescence intensity within the 3D volume which was quantified via ImageJ. Imaging for the 2D and 3D assays was performed using identical laser power and detector gain settings.

### 4.10. 2D Optical Assay via Silver Metallization

To establish a baseline for the optical readout, an enzymatically amplified silver metallization assay was performed on a 2D surface. This model assay utilized the same biotin-streptavidin interaction described in the fluorescence studies but substituted the fluorescent tag for an enzymatic (HRP) reporter. The initial functionalization, blocking, and washing steps to create the streptavidin detection spots were identical to the 2D fluorescence assay protocol. Following the preparation of the dual-layer PDMS assembly, serial dilutions of Strep-HRP were prepared in 1 mg mL^-1^ BSA in 0.05% PBST, with concentrations ranging from 2.5 pM to 2000 pM. A 100 μL volume of each dilution was added to the reservoir wells (*n* = 3 per condition) and incubated for 2 h on a plate shaker at 60 rpm.

After incubation and subsequent washing, the slides were rinsed with DI water to remove residual salts and dried. Equal volumes (33 μL) of EnzMet components A, B, and C were sequentially added to the wells and incubated for 4, 4 and 30 min, respectively to initiate silver metallization. The reaction was terminated by dipping the slides in DI water, followed by drying. The resulting darkness at the detection spots due to metal deposition was quantified by scanning the slides with a standard office scanner (MFC-L2710DW, Brother, USA). The optical darkness was quantified as the mean grayscale intensity of the detection spot using ImageJ.

### 4.11. 3D Optical Assay via Silver Metallization

To evaluate the sensitivity enhancement provided by the 3D hydrogel matrix for silver metallization-based readouts the experimental setup was identical to the 3D fluorescence assay protocol. Following overnight equilibration, the detection spots were incubated for 2 h with serial dilutions of Strep-HRP (*n* = 3 per condition) with concentrations ranging from 2.5 pM to 250 pM and washed. An additional wash with DI water was included to ensure the complete removal of residual salts from the hydrogel matrix. Enzymatic silver metallization was performed as detailed previously. The reaction was terminated by dipping the slide in DI water and the hydrogels were allowed to dry at RT. The imaging and quantification of the deposited silver was done as detailed previously.

### 4.12. Fabrication and Passivation of IDE Sensors

To enable electronic quantification of the silver metallization readout, flexible interdigitated electrodes (FIDE) were utilized as the transducer. The FIDEs consisted of a 10 mm x 10 mm pad size with a finger spacing of 0.254 mm. To define a localized sensing area, the FIDEs were passivated using polyimide tape, leaving a 2.5 mm diameter aperture in the center of the electrode array. A 6 mm diameter PDMS reservoir well was aligned over the passivated sensing region and secured with double-sided adhesive tape to contain the assay reagents **(Figure S3)**.

### 4.13. Electronic Impedance Spectroscopy (EIS) Assay via Silver Metallization

This assay utilized the same biotin-streptavidin model interaction as described previously. For the 2D baseline, the exposed IDE region was functionalized with 3 μL of 2.5 mg mL^-1^ Biotin-BSA for 1 h in a humidified chamber. Following functionalization, the sensors were blocked with 1% BSA and washed. For the 3D Bio-gel sensor, 1 μL of the Bio-gel precursor was pipetted onto the exposed FIDE region, photo-crosslinked, and swelled overnight in PBS to reach equilibrium.

Both the 2D and 3D Bio-gel sensors were incubated with 100 μL of Strep-HRP (3 nM) for 2 h. After washing to remove unbound streptavidin and residual salt content, the wells were refilled with 100 μL of fresh DI water. Baseline impedance spectrum measurements were recorded at 100 mV over a frequency range of 100 Hz to 1 MHz using an LCR meter (ZM2376, NF Corporation, Japan). Data acquisition was performed using a custom MATLAB script designed to perform automated electrochemical impedance spectroscopy (EIS). Following the baseline measurement, enzymatically amplified silver metallization was performed as detailed previously. The reaction was terminated by dipping the sensors in DI water. After subsequent washes with DI water to remove free ions from the sensing region, the wells were refilled with 100 μL of fresh DI water and the post-metallization EIS spectrum was recorded.

### 4.14. 3D COVID-19 Immunoassay for Antigen Specific Antibody Detection

SARS-CoV-2 nucleocapsid (NC-gels) and spike protein (SP-gels) functionalized hydrogels were used for the detection of antigen-specific antibodies (IgG and IgA). Hydrogel spots were formed by pipetting 1 μL of the precursor solutions into the wells of a glass-bottom 96-well plate followed by photo-crosslinking. The resulting detection spots were incubated in 1% BSA in 0.1% PBST overnight to reach equilibrium swelling and simultaneously block the matrix against non-specific protein adsorption. Post washing, 100 μL of clinical serum samples diluted in 1 mg mL^-1^ BSA in 0.05% PBST were added to each well and incubated for 2 h. The washing protocol was then repeated to remove unbound serum components. For single-analyte IgG detection, 100 μL of goat anti-human IgG-AF647 (100 μg mL^-1^) was added as the probe and incubated for 2 h, followed by a final washing cycle. Characterization of the 3D spots was performed via fluorescence microscopy as detailed previously to quantify the bound antibody signal.

To investigate the capacity for multiplexed detection within a single detection spot, SP-gels were evaluated for simultaneous IgG and IgA capture. In the probe addition step, 60 μL volumes of either goat anti-human IgG-PE (3.33 μg mL^-1^), goat anti-human IgA-AF647 (3.33 μg mL^-1^), or a combined cocktail of both probes were added. Following incubation and washing, the IgG and IgA signals were imaged via fluorescence microscopy using 561 nm and 640 nm laser lines, respectively, to verify target specificity and resolve distinct antibody signals.

### 4.15. ELISA

A high-binding 96-well plate was functionalized with SARS-CoV-2 nucleocapsid protein (2 μg mL^-1^ in PBS) via overnight incubation at 4°C (50 μL per well). After washing thrice with PBS, the wells were blocked with 1% BSA in 0.1% PBST for 1 hr at RT to minimize non-specific adsorption, followed by three additional PBS washes. Clinical serum samples diluted in 0.1% BSA and 0.05% PBST were introduced at 100 μL per well and allowed to incubate for 1 h. Following three washes with 0.05% PBST, 100 μL of mouse anti-human IgG Fc-HRP (1:1500 dilution) was added to each well and incubated for 1 h at 60 rpm. The plates were subsequently washed four times with 0.05% PBST before the addition of 50 μL TMB substrate. After 15 min, the enzymatic reaction was quenched with 50 μL of 1 M sulfuric acid. The optical density of each well was measured at 450 nm using a microplate reader (BioTek Synergy H4, Agilent, USA).

### 4.16. Fabrication of Lateral Flow Dipsticks

Lateral flow dipsticks were fabricated using laser-cut nitrocellulose (NC) membranes and absorbent pads. NC membranes were cut into 1 cm x 4.5 cm strips with one tapered end, while absorbent pads were cut into 1.2 cm x 5 cm strips. To assemble the dipsticks, the non-tapered end of the NC strip was sandwiched between two absorbent pads with a 0.5 cm overlap and secured with adhesive tape. Detection spots were localized 2 cm from the tapered end by carefully pipetting 0.5 μL of either Protein A/G (0.4 mg mL^-1^) for the traditional LFA (T-LFA) or 0.5 μL of AG-gel precursor (0.4 mg mL^-1^) for the hydrogel-enhanced LFA (H-LFA) which spreads to a diameter of approximately 2 mm upon contact with the NC membrane. Notably, while previous formulations for microplate-based assays utilized 6 wt% polymer concentrations, the AG-gel was optimized at 3 wt% to prevent the occlusion of the nitrocellulose pores. For testing multiplexability, we patterned a 2 x 4 array of alternating Protein A/G and non-functionalized spots either via functionalized hydrogels or direct protein addition. Immediately following deposition, HP-LFAs were photo-crosslinked for 3 min to immobilize the hydrogel matrix in situ. Both T-LFA and H-LFA dipsticks were dried for 2 h at 37°C, sealed and store at 4°C for use within two weeks.

### 4.17. Fluorescence based Lateral Flow Assay

To evaluate the comparative sensitivity of the HP-LFA and P-LFA platforms, an IgG capture assay was conducted using serial dilutions of goat anti-human IgG-AF647 ranging from 1 pM to 2500 pM (*n* = 3 per condition). Dilutions were prepared in 1% BSA in 0.1% PBST to minimize non-specific interaction with the NC fibers. Dipsticks were immersed in wells containing 500 μL of the probe solution for 30 min to allow for complete capillary uptake and analyte binding. The setup for performing this assay is shown in **Figure S4** For the multiplexed LFAs, goat anti-human IgG-AF647 at 2500 pM was used as the sample. Following a 30 min drying period at RT, the NC membranes were mounted onto standard microscope glass slides. The captured IgG was quantified via fluorescence microscopy by orienting the membrane-side toward the laser and the mean fluorescence intensities were determined using ImageJ.

### 4.18. Statistical Analysis

All data are presented as the mean ± standard deviation (SD) unless otherwise specified. Statistical analyses were performed using GraphPad Prism (Version 10). Differences among various hydrogel formulations were evaluated using t-tests, one-way or two-way ANOVAs followed by Tukey’s multiple comparisons test, with significance defined as p<0.05. Experimental replicates were maintained at n=3 to 5 per condition. The analytical LOD was calculated according to the three-sigma method as the concentration corresponding to the mean signal of the blank (PBS) plus three times its standard deviation.

## Acknowledgements

This work was supported by funding from the National Institute of Health (NIH) (GR00027857). The authors gratefully acknowledge the Optical Microscopy Core at Georgia Institute of Technology for providing access to imaging equipment.

## Data Availability Statement

The data that support the findings of this study are available from the corresponding author upon reasonable request.

## References

1 Wild, D. The immunoassay handbook: theory and applications of ligand binding, ELISA and related techniques. (Newnes, 2013).

2 Huang, J. et al. Explore how immobilization strategies affected immunosensor performance by comparing four methods for antibody immobilization on electrode surfaces. Scientific Reports 2022 12:1 12 (2022-12-23). 10.1038/s41598-022-26768-w

3 Susini, V. et al. Antibody-Antigen Binding Events: The Effects of Antibody Orientation and Antigen Properties on the Immunoassay Sensitivity | IntechOpen. Rapid Antigen Testing (2023/05/16). 10.5772/intechopen.1001374

4 Wilson, B. D. & Soh, H. T. Re-Evaluating the Conventional Wisdom about Binding Assays. Trends in Biochemical Sciences 45 (2020/08/01). 10.1016/j.tibs.2020.04.005

5 Hariri, A. A. et al. Improved immunoassay sensitivity and specificity using single-molecule colocalization. Nature Communications 2022 13:1 13 (2022-09-12). 10.1038/s41467-022-32796-x

6 Quartararo, A. J. et al. Ultra-large chemical libraries for the discovery of high-affinity peptide binders. Nature Communications 2020 11:1 11 (2020-06-23). 10.1038/s41467-020-16920-3

7 Gao, S., Guisán, J. M. & Rocha-Martin, J. Oriented immobilization of antibodies onto sensing platforms - A critical review. Analytica Chimica Acta 1189 (2022/01/02). 10.1016/j.aca.2021.338907

8 Zhang, Y. et al. Revolutionizing the capture efficiency of ultrasensitive digital ELISA via an antibody oriented-immobilization strategy. Journal of Materials Chemistry B 12 (2024/10/09). 10.1039/D4TB01141D

9 Belfakir, S. B. et al. Leveraging cellulose-binding domains to orient and immobilize single-domain antibodies onto paper-based immunoassays. Sensors and Actuators B: Chemical 439 (2025/09/15). 10.1016/j.snb.2025.137833

10 Sonny S. Mark, †, Neelakantapillai Sandhyarani, §, Changcheng Zhu, Christine Campagnolo, a. & Batt‡, C. A. Dendrimer-Functionalized Self-Assembled Monolayers as a Surface Plasmon Resonance Sensor Surface. Langmuir 20 (June 29, 2004). 10.1021/la0495276

11. Herrmann, A. Surface-Bound Functional Hydrogel Networks Ph.D. thesis, Freie Universität, (2020).

12 Tavakoli, J. & Tang, Y. Hydrogel Based Sensors for Biomedical Applications: An Updated Review. Polymers 9 (2017 Aug 16). 10.3390/polym9080364

13 Herrmann, A., Haag, R. & Schedler, U. Hydrogels and Their Role in Biosensing Applications. Advanced Healthcare Materials 10, 2100062 (2021). 10.1002/adhm.202100062

14 Jung, I. Y., Kim, J. S., Choi, B. R., Lee, K. & Lee, H. Hydrogel Based Biosensors for In Vitro Diagnostics of Biochemicals, Proteins, and Genes. Advanced Healthcare Materials 6, 1601475 (2017). 10.1002/adhm.201601475

15 Rubina, A. Y. et al. Hydrogel-Based Protein Microchips: Manufacturing, Properties, and Applications. BioTechniques 34 (2003-5-1). 10.2144/03345rr01

16 Herrmann, A., Kaufmann, L., Dey, P., Haag, R. & Schedler, U. Bioorthogonal in Situ Hydrogels Based on Polyether Polyols for New Biosensor Materials with High Sensitivity. ACS applied materials & interfaces. 10, 11382–11390 (2018). 10.1021/acsami.8b01860

17 Krage, C. et al. Three-Dimensional Polyglycerol–PEG-Based Hydrogels as a Universal High-Sensitivity Platform for SPR Analysis. Analytical Chemistry 97 (March 13, 2025). 10.1021/acs.analchem.5c00499

18 NW, C., et al. Multiplexed detection of mRNA using porosity-tuned hydrogel microparticles - PubMed. Analytical chemistry 84 (11/06/2012). 10.1021/ac302128

19 Kim, H. J. et al. Highly sensitive three-dimensional interdigitated microelectrode biosensors embedded with porosity tunable hydrogel for detecting proteins. Sensors and actuators. 302, 127190 (2020). 10.1016/j.snb.2019.127190

20 Chan, D. et al. Combinatorial Polyacrylamide Hydrogels for Preventing Biofouling on Implantable Biosensors. Advanced Materials 34 (2022/06/01). 10.1002/adma.202109764

21 Zhu, J. Bioactive modification of poly(ethylene glycol) hydrogels for tissue engineering. Biomaterials 31 (2010/06/01). 10.1016/j.biomaterials.2010.02.044

22 Randriantsilefisoa, R. et al. Highly sensitive detection of antibodies in a soft bioactive three-dimensional bioorthogonal hydrogel. Journal of Materials Chemistry B 7, 3220–3231 (2019). 10.1039/c9tb00234k

23 Luo, Q. et al. Portable functional hydrogels based on silver metallization for visual monitoring of fish freshness. Food Control 123 (2021/05/01). 10.1016/j.foodcont.2020.107824

24 Shohatee, D., Keifer, J., Schimmel, N., Mohanty, S. & Ghosh, G. Hydrogel-based suspension array for biomarker detection using horseradish peroxidase-mediated silver precipitation. Analytica chimica acta : An International Journal Devoted to All Branches of Analytical Chemistry 999, 132–138 (2018). 10.1016/j.aca.2017.10.033

25 Choi, J. R. et al. Lateral Flow Assay Based on Paper–Hydrogel Hybrid Material for Sensitive Point-of-Care Detection of Dengue Virus. Advanced Healthcare Materials 6 (2017/01/01). 10.1002/adhm.201600920

26 Mahardika, I. H. et al. From Pregnancy to Pathogens: Boosting Lateral Flow Assays Sensitivity with a Hydrogel Reaction Trap. Advanced Materials Interfaces 11 (2024/09/01). 10.1002/admi.202400341

27 Fairbanks, B. D. et al. A Versatile Synthetic Extracellular Matrix Mimic via Thiol-Norbornene Photopolymerization. Advanced Materials 21 (2009/12/28). 10.1002/adma.200901808

28 Mora-Boza, A. et al. Facile Photopatterning of Perfusable Microchannels in Synthetic Hydrogels to Recreate Microphysiological Environments. Advanced Materials 35 (2023). 10.1002/adma.202306765

29 Charles, P. T. et al. Fabrication and characterization of 3D hydrogel microarrays to measure antigenicity and antibody functionality for biosensor applications. Biosensors & bioelectronics. 20, 753–764 (2004). 10.1016/j.bios.2004.04.007

30 Son, K. H., Lee, J. W., Son, K. H. & Lee, J. W. Synthesis and Characterization of Poly(Ethylene Glycol) Based Thermo-Responsive Hydrogels for Cell Sheet Engineering. Materials 2016, Vol. 9, 9 (2016-10-20). 10.3390/ma9100854

31 Parlato, D. M., Reichert, D. S., Barney, D. N. & Murphy, D. W. L. Poly(ethylene glycol) Hydrogels with Adaptable Mechanical and Degradation Properties for Use in Biomedical Applications. Macromolecular bioscience 14 (2014 Jan 25). 10.1002/mabi.201300418

32 Peppas, N. A. & Merrill, E. W. Crosslinked poly(vinyl alcohol) hydrogels as swollen elastic networks. Journal of Applied Polymer Science 21 (1977/07/01). 10.1002/app.1977.070210704

33 Silverton, E. W. et al. Three-dimensional structure of an intact human immunoglobulin. Proceedings of the National Academy of Sciences 74 (1977-11). 10.1073/pnas.74.11.5140

34 Granath, K. A. & Kvist, B. E. Molecular weight distribution analysis by gel chromatography on sephadex. Journal of Chromatography A 28 (1967/01/01). 10.1016/S0021-9673(01)85930-6

35 Rafat, N. et al. Inexpensive High-Throughput Multiplexed Biomarker Detection Using Enzymatic Metallization with Cellphone-Based Computer Vision. ACS Sensors 8 (February 8, 2023). 10.1021/acssensors.2c01429

36 Armbruster, D. A. & Pry, T. Limit of blank, limit of detection and limit of quantitation. Clin Biochem Rev 29 Suppl 1, S49–52 (2008).

37 Ali, S. M. et al. Microscale Multiplexed Antigen-Specific Antibody Fc Profiling for Point-of-Care Diagnosis of Tuberculosis. medRxiv, 2025.2011.2015.25340307 (2025). 10.1101/2025.11.15.25340307

38 Elkhiyari, K. et al. Inexpensive High-Throughput Multiplexed Cytokine Detection for Tuberculosis Diagnostics Using Amplified Enzymatic Metallization. bioRxiv, 2026.2001.2027.700981 (2026). 10.64898/2026.01.27.700981

39 Senthil, M. et al. Multiplexed High-Throughput Detection of Schistosomiasis Biomarkers Using Enhanced Nanoscale Enzymatic Metallization. ACS Nanoscience Au (2026). 10.1021/acsnanoscienceau.6c00013

40 Rafat, N. et al. Enhanced Enzymatically Amplified Metallization on Nanostructured Surfaces for Multiplexed Point-of-Care Electrical Detection of COVID-19 Biomarkers. Small 18 (2022/12/01). 10.1002/smll.202203309

41 Yang, L. et al. Design considerations in the use of interdigitated microsensor electrode arrays (IMEs) for impedimetric characterization of biomimetic hydrogels. Biomedical Microdevices 2010 13:2 13 (2010-11-23). 10.1007/s10544-010-9492-4

42 Hanhao Zhang et al. High throughput electronic detection of biomarkers using enzymatically amplified metallization on nanostructured surfaces. Analytical Methods 16 (2024/11/28). 10.1039/D4AY01657B

43 Peddireddy, S. P. et al. Antibodies targeting conserved non-canonical antigens and endemic coronaviruses associate with favorable outcomes in severe COVID-19. Cell Rep 39, 111020 (2022). 10.1016/j.celrep.2022.111020

44 Jazayeri, M. H., Pourfathollah, A. A., Rasaee, M. J., Porpak, Z. & Jafari, M. E. The concentration of total serum IgG and IgM in sera of healthy individuals varies at different age intervals. Biomedicine & Aging Pathology 3 (2013/10/01). 10.1016/j.biomag.2013.09.002

45 Shirshahi, V. & Liu, G. Enhancing the analytical performance of paper lateral flow assays: From chemistry to engineering. TrAC Trends in Analytical Chemistry 136 (2021/03/01). 10.1016/j.trac.2021.116200

46 Weimer, B. C., Walsh, M. K. & Wang, X. Influence of a poly-ethylene glycol spacer on antigen capture by immobilized antibodies. Journal of Biochemical and Biophysical Methods 45 (2000/09/11). 10.1016/S0165-022X(00)00114-7

47 Charlermroj, R. et al. Development of a microarray lateral flow strip test using a luminescent organic compound for multiplex detection of five mycotoxins. Talanta 233 (2021/10/01). 10.1016/j.talanta.2021.122540

48 Gantelius, J. et al. A Lateral Flow Protein Microarray for Rapid and Sensitive Antibody Assays. International Journal of Molecular Sciences 2011, Vol. 12, Pages 7748-7759 12 (2011-11-09). 10.3390/ijms12117748

49 Lee, K. W. et al. Instrumentation-Free Semiquantitative Immunoanalysis Using a Specially Patterned Lateral Flow Assay Device. Biosensors 2020, Vol. 10, Page 87 10 (2020-07-31). 10.3390/bios10080087

50 Hong, S. Y., Park, Y. M., Jang, Y. H., Min, B.-H. & Yoon, H. C. Quantitative lateral-flow immunoassay for the assessment of the cartilage oligomeric matrix protein as a marker of osteoarthritis. BioChip Journal 6, 213–220 (2012). 10.1007/s13206-012-6303-4

51 Gantelius, J. et al. A lateral flow protein microarray for rapid determination of contagious bovine pleuropneumonia status in bovine serum. Journal of Microbiological Methods 82 (2010/07/01). 10.1016/j.mimet.2010.03.007

52 Tisone, B. O. F. C. Lateral flow assays using two dimensional features. (2012).

53 Zhang, H. et al. Sample-Sparing Multiplexed Antibody Fc Biomarker Discovery Using a Reconfigurable Integrated Microfluidic Platform. Lab on a Chip (2025). 10.1039/D5LC00042D

54 Saha, A. et al. Deep humoral profiling coupled to interpretable machine learning unveils diagnostic markers and pathophysiology of schistosomiasis. Sci Transl Med 16, eadk7832 (2024). 10.1126/scitranslmed.adk7832

55 Ramachandran, S. et al. A Rapid, Multiplexed, High-Throughput Flow-Through Membrane Immunoassay: A Convenient Alternative to ELISA. Diagnostics (Basel) 3, 244–260 (2013). 10.3390/diagnostics3020244

56 Wu, Y. et al. Quantitative assessment of a novel flow-through porous microarray for the rapid analysis of gene expression profiles. Nucleic Acids Res 32, e123 (2004). 10.1093/nar/gnh118

57 Joung, H. A. et al. Paper-based multiplexed vertical flow assay for point-of-care testing. Lab Chip 19, 1027–1034 (2019). 10.1039/c9lc00011a

58 Goryacheva, I. Y. in Comprehensive Analytical Chemistry Vol. 72 (ed Irina Yu Goryacheva) 133-161 (Elsevier, 2016).

59 Shakeri, A. et al. Noncontact 3D Bioprinting of Proteinaceous Microarrays for Highly Sensitive Immunofluorescence Detection within Clinical Samples. ACS nano. 18, 31506–31523 (2024). 10.1021/acsnano.4c12460

60 Choi, G. et al. Portable microfluidic immunoassay platform for the detection of inflammatory protein biomarkers. Sensors & Diagnostics 3 (2024/04/18). 10.1039/D3SD00258F

61 Han, G.-R. et al. Deep Learning-Enhanced Paper-Based Vertical Flow Assay for High-Sensitivity Troponin Detection Using Nanoparticle Amplification. ACS Nano 18 (October 4, 2024). 10.1021/acsnano.4c05153

62 Rallapalli, Y. et al. Electronic Detection of Functional Cellular Immunity Using Enzymatic Metallization. ACS Omega (2026). 10.1021/acsomega.5c11489

63 Rudge, J. et al. Electronic Immunoassay Using Enzymatic Metallization on Microparticles. ACS Omega (2023). 10.1021/acsomega.3c01939

64 Canal, T. & Peppas, N. A. Correlation between mesh size and equilibrium degree of swelling of polymeric networks. Journal of Biomedical Materials Research 23 (1989/10/01). 10.1002/jbm.820231007

65 EW, M., KA, D. & C, S. Partitioning and diffusion of solutes in hydrogels of poly(ethylene oxide) - PubMed. Biomaterials 14 (1993 Dec). 10.1016/0142-9612(93)90154-t

