## Supplementary Information for "Multimodal Sensitivity Enhancement of Binding-based Diagnostic Assays Using Functional Hydrogels"

**Table S1. Hydrogel formulations and component quantities required for 100 µL batch volumes.**

| Formulation Designation | Degree of Crosslinking (%) | PEG-4aNB (mg) | Crosslinker MW | SH-PEG-SH (mg) | SH-PEG-Biotin (mg) | PEG 600 Porogen (µL) |
|---|---|---|---|---|---|---|
| **10/100 NF-gel** | 100% | 10.0 | 5 kDa | 5.0 | 0.0 | 0.0 |
| **10/87.5 Bio-gel** | 87.5% | 10.0 | 5 kDa | 4.375 | 0.25 | 0.0 |
| **10/75 Bio-gel** | 75% | 10.0 | 5 kDa | 3.75 | 0.5 | 0.0 |
| **10/50 Bio-gel** | 50% | 10.0 | 5 kDa | 2.5 | 1.0 | 0.0 |
| **6/100 NF-gel** | 100% | 6.0 | 5 kDa | 3.0 | 0.0 | 0.0 |
| **6/87.5 Bio-gel** | 87.5% | 6.0 | 5 kDa | 2.625 | 0.15 | 0.0 |
| **6/75 Bio-gel** | 75% | 6.0 | 5 kDa | 2.25 | 0.3 | 0.0 |
| **6/50 Bio-gel** | **50%** | **6.0** | **5 kDa** | **1.5** | **0.6** | **0.0** |
| **6/50 Bio-gel (1 kDa)** | 50% | 6.0 | 1 kDa | 0.3 | 0.6 | 0.0 |
| **6/50 Bio-gel (314 Da)** | 50% | 6.0 | 314 Da | 0.0942 | 0.6 | 0.0 |
| **6/50 Bio-gel (porogen)** | 50% | 6.0 | 5 kDa | 1.5 | 0.6 | 20.0 |

**Note: All components were dissolved in a working buffer composed of 10 mM HEPES in phosphate buffered saline (PBS). For photo-initiated crosslinking, a stock solution of LAP was added to each mixture such that the final concentration of LAP in all 100 µL precursor formulations was 1 mM. The highlighted optimal formulation was used for all sensitivity testing.*

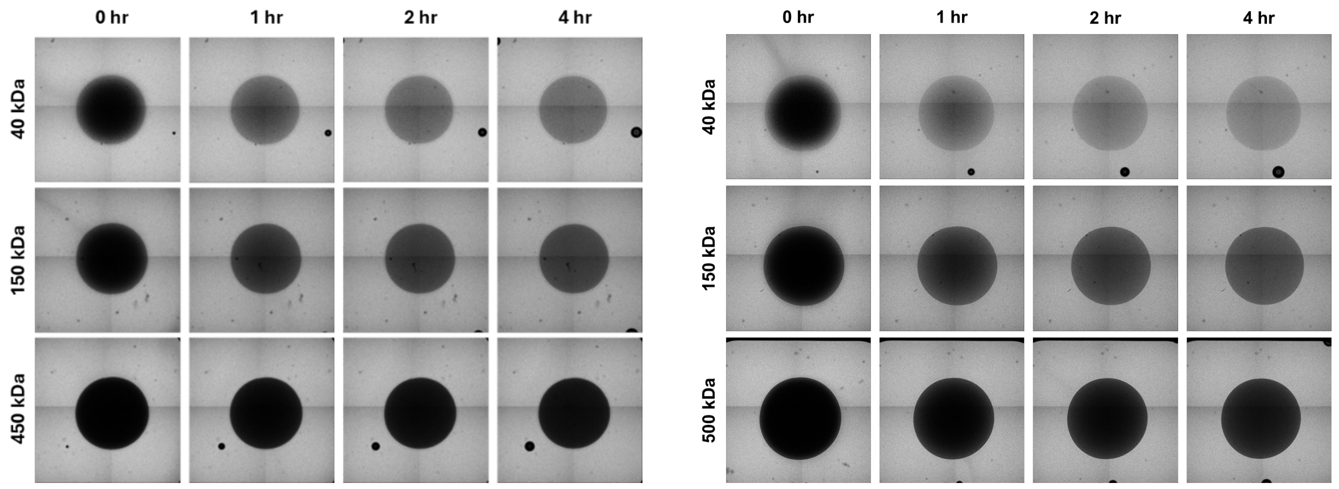


A. Without Porogen B. With Porogen

**Figure S1. Time series diffusion profile of FITC tagged dextran with varying molecular weights into Bio-gel detection spots (A)** with and **(B)** without porogen modification

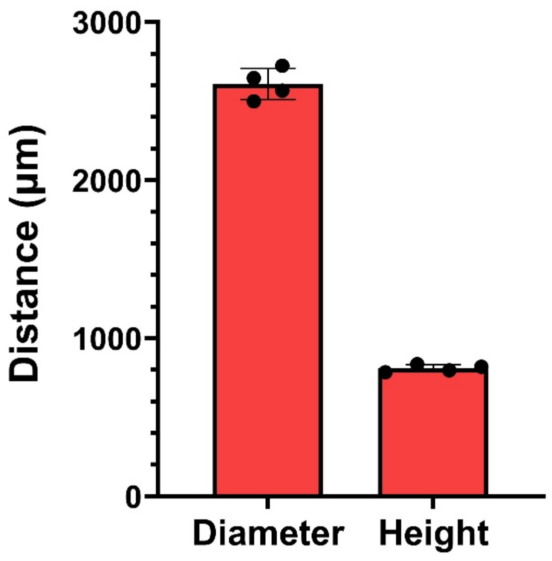


**Figure S2. Mean equilibrium diameter and height of a 1 $\mu$L Bio-gel sensing spot**

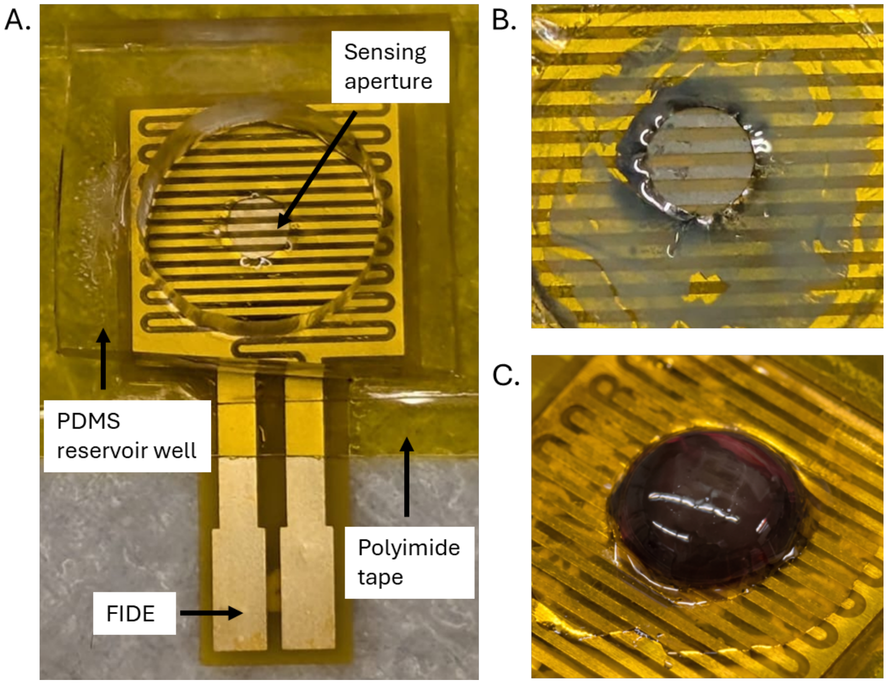


**Figure S3. Assembly of the flexible interdigitated electrode (FIDE) sensors and visual evaluation of silver metallization. (A)** Image of assembled FIDE platform used for Electronic Impedance Spectroscopy (EIS). **(B)** Image of the unmodified 2D electrode surface following the silver metallization assay. **(C)** Image of the 3D Bio-gel on the FIDE surface following the silver metallization assay

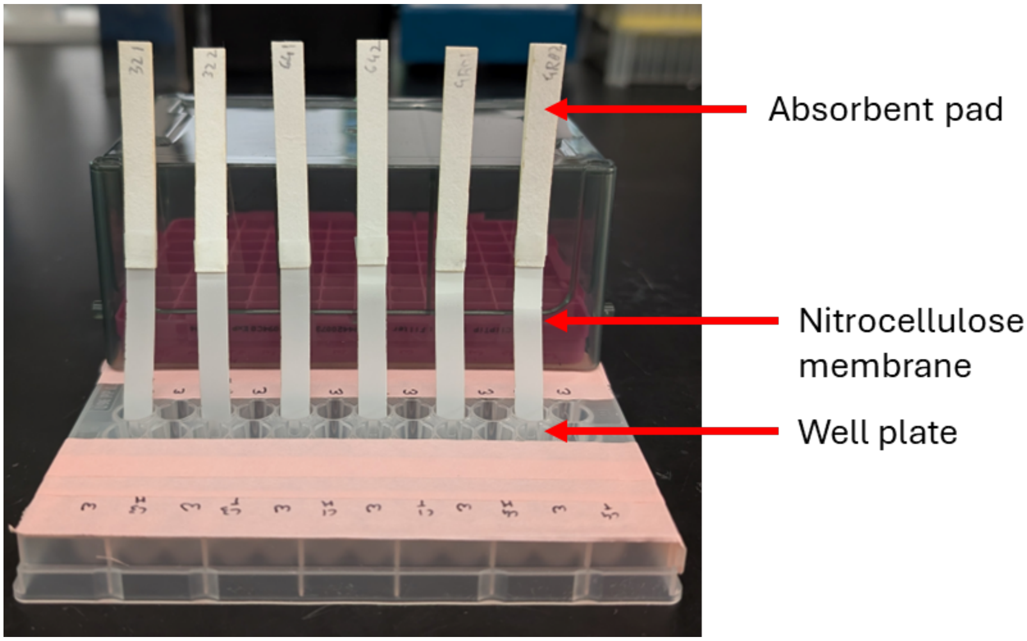


**Figure S4. Image of the setup for performing binding assays on LFA dipsticks**
